# Fructose-2,6-bisphosphate restores DNA repair activity of PNKP and ameliorates neurodegenerative symptoms in Huntington’s disease

**DOI:** 10.1101/2023.10.26.564220

**Authors:** Anirban Chakraborty, Sravan Gopalkrishnashetty Sreenivasmurthy, Wyatt Miller, Weihan Huai, Tapan Biswas, Santi Mohan Mandal, Lisardo Boscá, Balaji Krishnan, Gourisankar Ghosh, Tapas Hazra

## Abstract

Huntington’s disease (HD) and spinocerebellar ataxia type 3 (SCA3) are the two most prevalent polyglutamine (polyQ) neurodegenerative diseases, caused by CAG (encoding glutamine) repeat expansion in the coding region of the huntingtin (HTT) and ataxin-3 (ATXN3) proteins, respectively. We have earlier reported that the activity, but not the protein level, of an essential DNA repair enzyme, polynucleotide kinase 3’-phosphatase (PNKP), is severely abrogated in both HD and SCA3 resulting in accumulation of double-strand breaks in patients’ brain genome. While investigating the mechanistic basis for the loss of PNKP activity and accumulation of DNA double-strand breaks leading to neuronal death, we observed that PNKP interacts with the nuclear isoform of 6-phosphofructo-2-kinase fructose-2,6-bisphosphatase 3 (PFKFB3). Depletion of PFKFB3 markedly abrogates PNKP activity without changing its protein level. Notably, the levels of both PFKFB3 and its product fructose-2,6 bisphosphate (F2,6BP), an allosteric modulator of glycolysis, are significantly lower in the nuclear extracts of post-mortem brain tissues of HD and SCA3 patients. Supplementation of F2,6BP restored PNKP activity in the nuclear extracts of patients’ brain. Moreover, intracellular delivery of F2,6BP restored both the activity of PNKP and the integrity of transcribed genome in neuronal cells derived from striatum of HD mouse. Importantly, supplementing F2,6BP rescued the HD phenotype in Drosophila, suggesting F2,6BP to serve in vivo as a cofactor for the proper functionality of PNKP and thereby, of brain health. Our results thus provide a compelling rationale for exploring the therapeutic use of F2,6BP and structurally related compounds for treating polyQ diseases.

**Significance:** To unravel the biological basis for the loss of PNKP activity in HD and SCA3, the two most prevalent polyglutamine neurodegenerative disorders, we analyzed PNKP interactome and found that the nuclear isoform of a glycolytic enzyme PFKFB3 associated with PNKP and other repair proteins forming a multiprotein complex. Surprisingly, we found that PFKFB3 and its biosynthetic product, F2,6BP are significantly low in the affected region of patients’ brain. Exogenous addition of F2,6BP restored PNKP activity in patients’ brain nuclear extract. Moreover, supplementing F2,6BP in HD cells and fruit flies restored genome integrity and rescued the disease symptoms. While there is no curative therapy for HD/SCA3, except symptom management, our discovery suggests that F2,6BP supplementation would be a promising therapeutic option.

## Introduction

Polyglutamine (polyQ) diseases are dominant, heritable neurodegenerative disorders that are manifested by progressive deterioration of cognitive and motor functions^1–6^. Huntington’s disease (HD) and Spinocerebellar ataxia type 3 (SCA3) are the most common polyglutamine (polyQ) diseases worldwide. HD is caused by a polyQ expansion (>36) in the N-terminal region of huntingtin (HTT) protein^1^. Likewise, SCA3 is attributed to unstable glutamine repeat expansion from 12-41 in healthy individuals to 62-84 in patients in the C-terminal region of ataxin-3 (ATXN3) protein^4^. Despite a well-defined genetic basis, the precise pathophysiological mechanisms underlying HD and SCA3 remain elusive^7,8^. Recent genome-wide association studies and genetic data indicate deficient DNA repair as a contributing factor in the pathogenesis of polyQ diseases^9,10^.

DNA strand breaks, a common occurrence under various physiological processes, or during DNA base excision repair (BER), lead to blocked DNA termini at the break sites that can impede DNA repair and stall elongating RNA polymerases^11^. 3’-phosphate (3’-P) is one of such major blocked DNA termini in mammalian cells. Polynucleotide kinase 3’-phosphatase (PNKP), a bifunctional DNA end-processing enzyme with 3’-phosphatase and 5’-kinase activities, is a major 3’-phosphatase in mammalian cells^12,13^ and thus, it participates in multiple DNA repair pathways, including BER/single-strand break (SSBR)^14,15^ and classical non-homologous end-joining (C-NHEJ)-mediated DSB repair^16–21^. The knockout (KO) of the gene encoding PNKP leads to post-natal death in mice due to a defect in neurogenesis and oligodendrogenesis^22^. Multiple mutations have been mapped to the *Pnkp* gene that are responsible for neurological disorders, including severe progressive polyneuropathy, cerebellar atrophy, microcephaly, mild epilepsy, seizures, developmental delay, and intellectual disability^23–25^.

Our previous research demonstrated the association of HTT and ATXN3 with PNKP, RNA polymerase II (RNAPII) and several other DNA repair proteins, forming a transcription-coupled DNA repair complex^19, 21, 26, 27^. We further found that the 3’-phosphatase activity of PNKP, but not the protein level, was severely compromised in the affected brain regions of HD and SCA3 mice and, in the postmortem brain tissues (cerebellum) of SCA3 patients. The inactivation of PNKP in oxidation sensitive neurons results in a progressive accumulation of DNA strand breaks that ultimately activate pro-apoptotic pathways causing neuronal death^26,27^. Intriguingly, our studies showed amelioration of neurotoxicity by overexpression of PNKP in cells expressing polyQ expanded mutant HTT (mHTT)^27^ and ATXN3 (mATXN3)^26^ and most importantly, in a functional assay using a Drosophila model of SCA3^19^.

In this study, we delved into understanding the mechanism behind the loss of PNKP activity in HD and SCA3. We identified 6-phosphofructo-2-kinase/fructose-2,6-bisphosphatase 3 (PFKFB3) as a new component of the transcription-coupled non-homologous end-joining (TC-NHEJ) repair complex. PFKFB3, one of the four homodimeric bifunctional enzymes (PFKFB1-4), can reversibly phosphorylate fructose-6-phosphate (F6P) to fructose-2,6-bisphosphate (F2,6BP)^28^. PFKFB3 is the only isoform that translocates to the nucleus and targeted disruption of the mouse PFKFB3 is embryonic lethal, indicating its important role in the nucleus^29,30^. Notably, we found that F2,6BP can restore 3’-phosphatase activity of PNKP in HD and SCA3 patients’ brain nuclear extracts *in vitro*. Delivering F2,6BP^31,32^ into HD-mouse-derived striatal neurons protected them by reinstating PNKP activity and maintaining transcribed genome integrity. Finally, supplementing F2,6BP in a Drosophila model system expressing human *Htt*128Q^33^ rescued the impaired motor function phenotype, providing a compelling rationale for exploring the therapeutic potential of the metabolite or its analog against such polyQ diseases.

## Results and Discussion

### PFKFB3 is a component of the TC-NHEJ complex in mammalian cells

We previously described that the DNA repair process in mammalian brains involves dynamic multi-protein complexes^19, 20, 21, 27^. A key finding in our current study, identified through 2D gel and mass spectrometric (MS) analysis, is the presence of the glycolysis modulator PFKFB3 within the TC-NHEJ complex **(Supplementary Fig. S1, Supplementary Table S1)**. Immunoprecipitation (IP) using anti-PFKFB3 or anti-DNA ligase IV (Lig IV; exclusively known for its role in C-NHEJ) antibodies (Abs) from the nuclear extracts of mice brains confirmed the presence of PFKFB3, ATXN3 and PNKP along with RNA polymerase II (RNAPII) and other known repair proteins in the TC-NHEJ complex **(Fig. 1A)**. We have earlier reported that HTT forms a complex with BER/SSBR proteins along with RNAPII, thereby implicating its role in transcription-coupled DNA repair^27^. Therefore, to further assess HTT’s role in TC-NHEJ, we performed co-immunoprecipitation using anti-PFKFB3 and anti-Lig IV Abs and indeed found HTT in both PFKFB3 and Lig IV immunocomplexes (ICs). A reverse immuno-pulldown using anti-HTT antibody further confirmed the association of HTT with PFKFB3, ATXN3, PNKP and RNAPII, implicating the role of HTT in TC-NHEJ **(Fig. 1B)**. The absence of homologous recombination (HR) protein, RAD51 in Lig IV (a C-NHEJ protein) IC provided specificity of the complex formation **(Fig. 1A, lane 5)**. However, RAD51 was found to be associated with PFKFB3 **(Fig. 1A, lane 4)**, which is consistent with a recent study reporting the role of PFKFB3 in DSB repair via the HR pathway^34^.

**Fig. 1:**
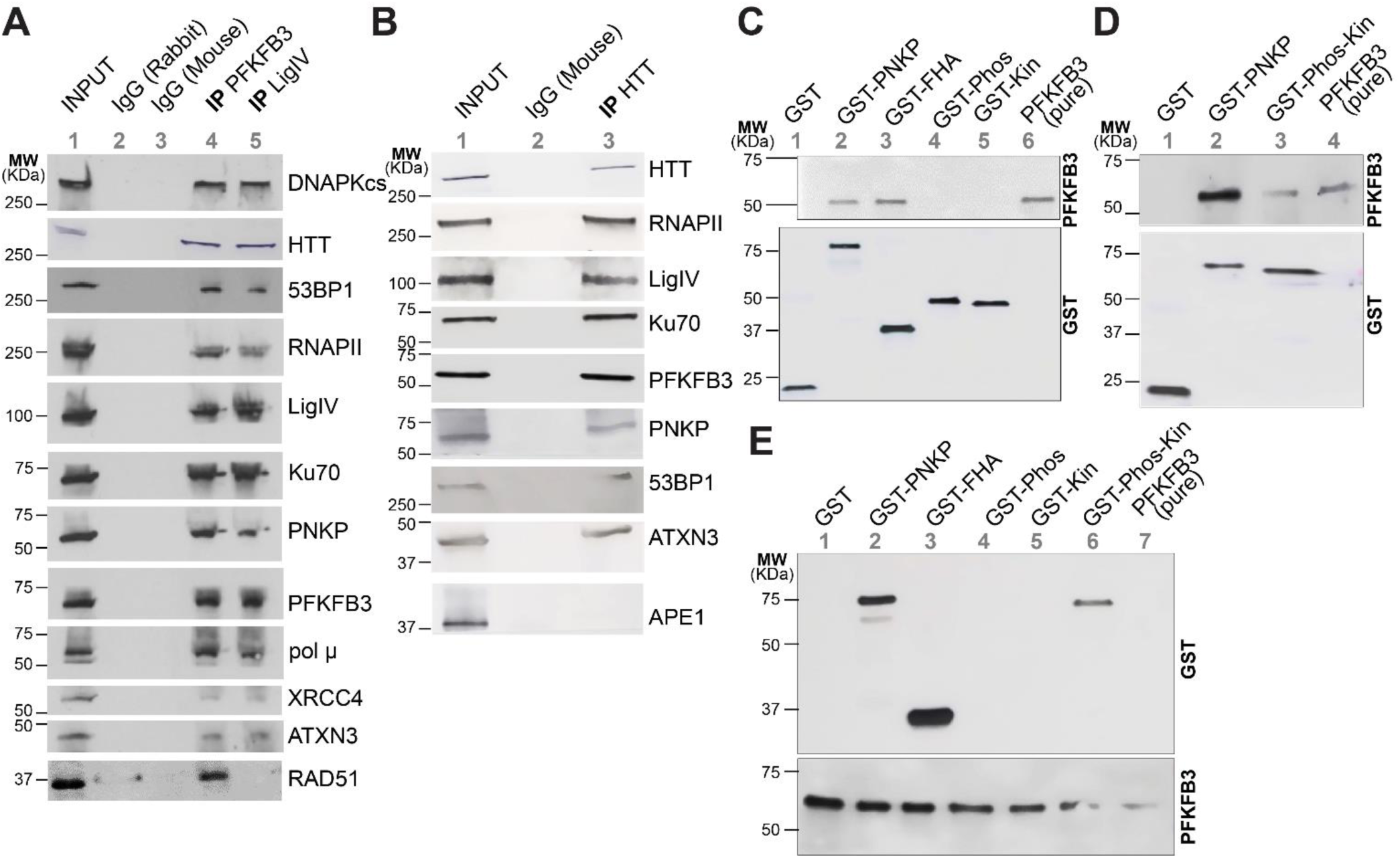
Partial characterization of TC-NHEJ complexes involving PNKP, PFKFB3 and HTT. **A.** Benzonase-treated nuclear extracts (NEs) from wild-type (WT) mouse (6 months old) brain were immunoprecipitated (IP’d) with anti-PFKFB3 (lane 4) and anti-Lig IV (lane 5) antibodies (Abs); or control IgG (Rabbit; lane 2 and mouse; lane 3) and tested for the presence of associated proteins (shown in the right) using specific Abs (n=3). **B.** Similar extract was IP’d with anti-HTT (lane 3) Ab or control mouse IgG (lane 2) and tested for the presence of associated proteins (shown in the right) using specific Abs (n=3). **C.** Full length WT PNKP and individual PNKP domains (FHA, phosphatase and kinase) were expressed as GST tagged proteins and allowed to bind to glutathione sepharose beads. **D.** Similar interaction with full length WT PNKP and fused (phosphatase + kinase) catalytic domain. Equimolar (20 pmol) amount of His-tagged PFKFB3 was subsequently bound to the full length PNKP/domains and after stringent wash, interaction was assessed by Western blot analysis with anti-PFKFB3 Ab **(Upper Panel)**. **Lower Panel**: Immunoblot shows bead bound GST/GST-PNKP or domains probed with anti-GST Ab. Lane 6 or lane 4: Purified PFKFB3 as control. For Fig. 1C, upper and lower panels were generated from two separate gels run in parallel. **E.** Full length WT PNKP and individual PNKP domains (FHA, phosphatase, kinase and phosphatase-Kinase) were expressed as GST tagged proteins and allowed to bind to His-tagged PFKFB3 (20 pmol) bound to cobalt-resins. After stringent wash, interaction was assessed by Western blot analysis with anti-GST Ab **(Upper Panel)**. **Lower Panel:** Immunoblot shows bead bound PFKFB3 with anti-PFKFB3 Ab. Lane 7: Purified PFKFB3 as control.

Next, we performed GST-pulldown experiments using purified recombinant full-length as well as different domains of PNKP to investigate if PNKP and PFKFB3 directly interact. The results revealed that the full-length and fork-head associated (FHA) domain of PNKP interacted directly with PNKP. Neither individual phosphatase nor kinase domain associated with PFKFB3; however, fused catalytic domain (phosphatase + kinase) showed modest but detectable association with PFKFB3 **(Fig. 1C**, 1D). Similar results were observed when we performed a reverse pulldown with His-tagged PFKFB3 **(Fig. 1E)**. Notably, it has been established previously that the FHA domain of PNKP directly interacts with other DNA repair proteins, namely XRCC1 (involved in SSBR)^35^ and XRCC4 (involved in C-NHEJ)^17^. Our study underscores the likelihood of the FHA domain acting as a pivotal protein-protein interaction hub in orchestrating the formation of the pathway-specific repair complexes. Additionally, these results corroborate with a recent report that shows while phosphorylated XRCC1 stimulates PNKP by binding to its FHA domain, non-phosphorylated XRCC1 stimulates PNKP by interacting with the PNKP catalytic domain^35^.

### PFKFB3 is involved in DSB repair of transcribed genes in mammalian cells

To assess the role of PFKFB3 in TC-NHEJ, we depleted PFKFB3 in HEK293 cells by specific siRNA **(Supplementary Fig. S2A)**, followed by treatment with bleomycin (Bleo), a DSB-inducing radio-mimetic drug, and monitored the repair at various time points (3, 6, 9, 12 h post Bleo treatment). Control cells demonstrated efficient repair of DNA strand-breaks within 6-9 h post Bleo treatment, as indicated by the resolution of γH2AX (a DSB marker). In contrast, PFKFB3-depleted cells exhibited impaired repair, as shown by unresolved γH2AX even at 12 h post Bleo treatment **(Supplementary Fig. S2A)**. We used a complementary, long amplicon (LA)-qPCR-based assay to quantitatively determine whether the DNA strand breaks persisted in PFKFB3-depleted cells. In this assay, a relative decrease in the PCR product of the long amplicon (∼ 8-10 kb) vs. the short amplicon (∼ 250 bp) in transcribed or non-transcribed genes reflects accumulation of DNA damage. Persistent DSB was monitored in representative transcribed (*HPRT*, *POLB* and *POLR2A*) vs. non-transcribed genes (*NanoG*, *Oct4* and *MyH2*). Control siRNA-treated cells demonstrated repair in all genes within 9 h following Bleo treatment, as indicated by efficient LA-PCR product formation. However, in PFKFB3-depleted cells, DSBs persisted specifically in transcribed genes **(Supplementary Fig. S2B)**. Interestingly, the non-transcribed genes showed significant repair within 6-9 h, similar to the control cells **(Supplementary Fig. S2C)**. These data highlight a critical role of PFKFB3 in DSB repair of transcribed genes via the TC-NHEJ pathway. Intriguingly, another recent report also revealed the role of a glycolytic enzyme, aldolase A in DSB repair^36^, suggesting metabolic and DNA repair pathways are closely linked.

### F2,6BP potentiates the 3’-phosphatase activity of PNKP *in vitro*

We have previously demonstrated the critical role of PNKP in the preferential repair of transcribed genes in mammalian cells^18^. Given the direct interaction between PNKP and PFKFB3 forming a TC-NHEJ complex and PFKFB3’s role in transcribed genome specific repair, we assessed the functional relationship of these proteins in DNA repair activities and their relevance to neurodegenerative pathology. We discovered that PNKP activity (schematically shown in **Supplementary Fig. S3A**) was compromised in PFKFB3-depleted **(Fig. 2A)** nuclear extracts from HEK293 cells **(Fig. 2B, lane 3 vs. lane 2)**. Complementation with recombinant PFKFB3 failed to restore PNKP activity **(Fig. 2B, lane 4 vs. lane 3)**. PFKFB3, known to shuttle to the nucleus unlike other isoforms^30^, is primarily responsible for F2,6BP synthesis since its kinase activity is ∼700-fold higher than the phosphatase activity^29,37^. We hypothesized that PNKP utilizes F2,6BP as a cofactor for its catalytic activity. Consistent with this hypothesis, PNKP activity was restored in PFKFB3-depleted nuclear extracts in a dose-dependent manner when supplemented with F6P and ATP along with recombinant PFKFB3 **(Fig. 2B, lanes 5, 6 vs. lane 3)**. Neither the F6P alone, nor a related metabolite, fructose-1,6-bisphosphate (F1,6BP) restored PNKP activity **(Fig. 2B, lanes 9 and 10**, respectively). Since PFKFB3, in the presence of F6P and ATP, is expected to produce F2,6BP, we surmised that F2,6BP itself can induce PNKP activity. To confirm this, we supplemented PFKFB3-depleted nuclear extracts with purified F2,6BP and indeed observed restoration of the activity of PNKP in a dose-dependent manner **(Fig. 2B, lanes 7, 8 vs. lane 3)**. We also observed that incubation of F2,6BP alone did not release any radiolabeled phosphate indicating that 3’-phosphate release activity observed in our assays is PNKP-specific **(Supplementary Fig. S3B)**. We further validated the dose dependent activation of purified PNKP by F2,6BP (**Fig. 2C, lane 3 vs. lanes 4-7**). In contrast, purified PFKFB3 **(Fig. 2C, lane 8)** or F6P **(Fig. 2C, lane 9)** had no discernable effect in activating PNKP. We next tested if F2,6BP affected the 5’-kinase activity of PNKP. Surprisingly, F2,6BP abrogated the kinase activity of PNKP in a dose-dependent manner **(Supplementary Fig. S3C, lane 2 vs. lanes 3-5)**. The relevance of such a contrarian effect of the metabolite on bifunctional activities of PNKP warrants further investigation.

**Fig. 2:**
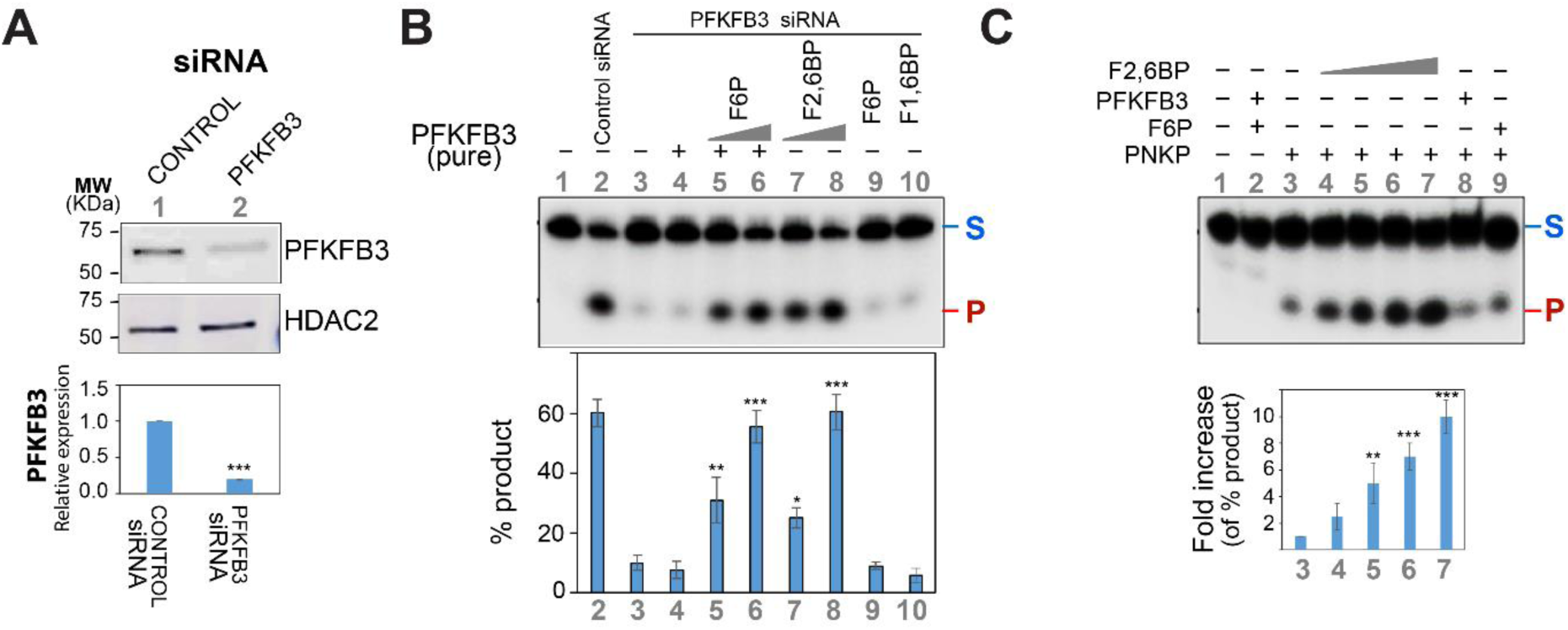
Effect of fructose-2,6-bisphosphate on 3’-phosphatase activity of PNKP. **A. Upper panel:** Western blot showing PFKFB3 and PNKP levels in the nuclear extracts of control vs. PFKFB3 siRNA transfected HEK293 cells. The **Lower panel** shows the quantitation of the relative PFKFB3 levels after normalization with nuclear loading control HDAC2. **B. Upper panel:** A ^32^P-labeled 3’-phosphate-containing oligo substrate (5 pmol) was incubated with the nuclear extract (250 ng) of control (lane 2) and PFKFB3-depleted HEK293 cells (lanes 3-10). Lane 4: purified PFKFB3 (200 fmol); lanes 5-6: increasing amounts of F6P (5 and 10 mM) were added along with purified PFKFB3 (200 fmol) and ATP (1 mM); lanes 7, 8: increasing amounts of F2,6BP (25 and 50 µM) were added; Lane 9: F6P (50 µM) and lane 10: F1,6BP (50 µM). Lane 1: No protein. **Lower panel:** Quantitation of the % released phosphate in the indicated lanes. S: Substrate and P: Released phosphate. **C. Upper panel:** 3’-phosphatase activity of purified PNKP alone (25 fmol, lane 3), PNKP plus increasing amounts of F2,6BP (5, 10, 15, and 20 µM, lanes 4-7), PNKP plus PFKFB3 (200 fmol, lane 8), PNKP plus F6P (25 µM, lane 9). Lane 1: substrate only; lane 2: PFKFB3 (200 fmol) plus F6P (25 µM). **Lower panel:** Quantitation of the fold change of the % released phosphate with the activity of purified PNKP considered arbitrarily as unity. S: Substrate and P: Released phosphate. In all the above cases, error bars show ±SD of the mean; n=3, *P<0.05; **P<0.01; ***P<0.005 (compared to lane 3 in **B** and **C**).

### PFKFB3 and F2,6BP levels are low in HD and SCA3 and exogenous F2,6BP restores PNKP activity

Our findings suggest that PNKP can utilize F2,6BP as a cofactor for its catalytic activity *in vitro*. We hypothesized that low PFKFB3 levels and/or activity might contribute to decreased PNKP activity and the accumulation of DNA damage in HD and SCA3 pathologies. To test this hypothesis, we first assessed the PNKP activity in the nuclear extracts of post-mortem HD patients’ brains (frontal cortex) vs. age/gender matched healthy controls. We observed a near complete abrogation of 3’-phosphatase activity of PNKP **(Supplementary Fig. S4)** consistent with previous observation in HD mice^27^. We further tested the F2,6BP level in the post-mortem HD patients that showed diminished PNKP activity, and the level was found to be about 2-fold decreased in the nuclear extracts of patients than in the corresponding controls **(Fig. 3A)**. The decreased level of F2,6BP correlated with a reduction in the PFKFB3 level in the respective HD patients (**Fig. 3B**). The level of PNKP protein remained comparable in HD patients and their age-matched controls **(Fig. 3B)**, consistent with our previous results^27^.

**Fig. 3:**
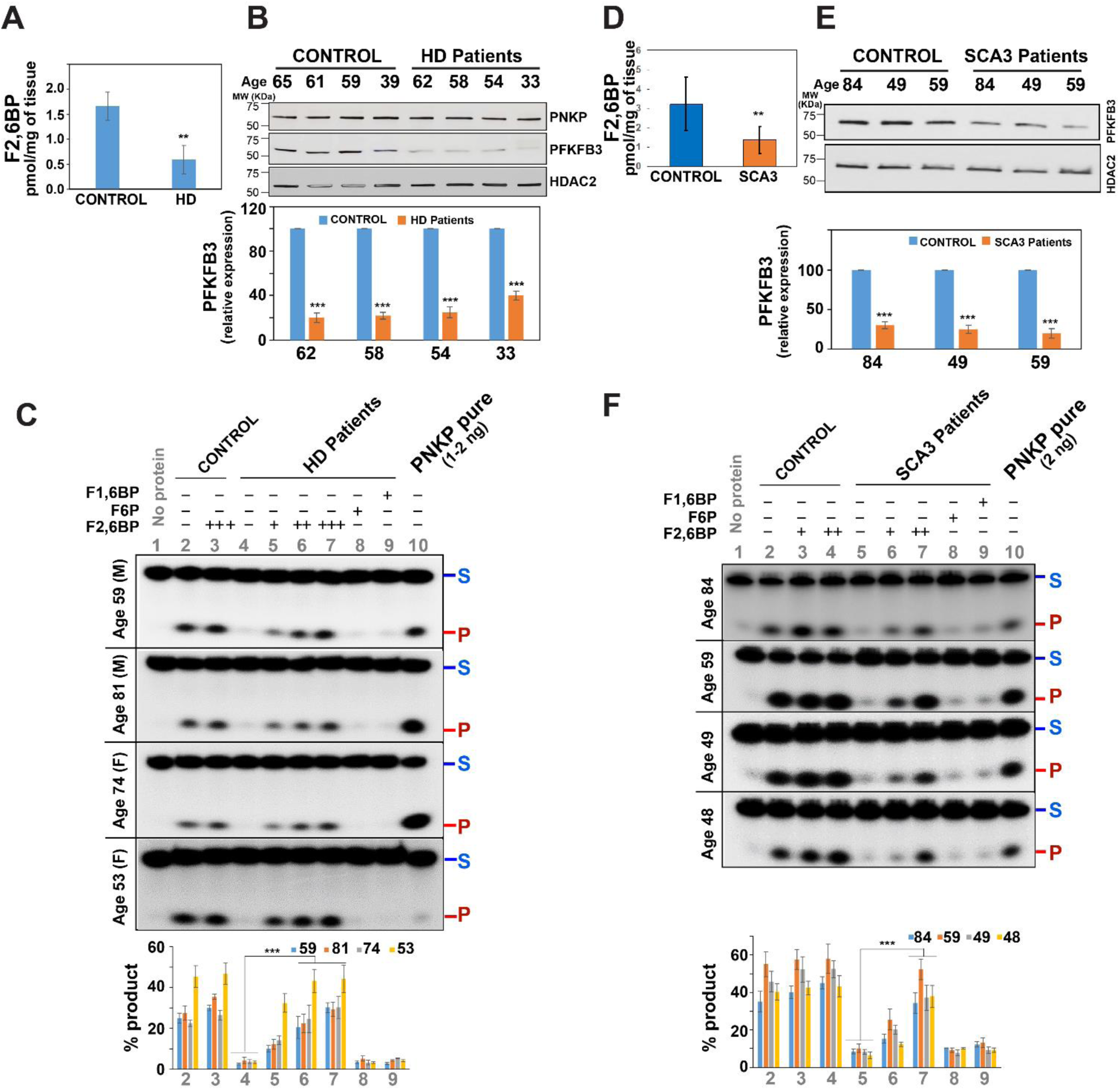
F2,6BP mediated restoration of the 3’-phosphatase activity of PNKP in the nuclear extracts of post-mortem HD and SCA3 patients’ brains. **A.** Bar diagram showing the relative levels of F2,6BP in the nuclear extract of control vs. HD patients’ frontal cortex (n=4). **B. Upper panel:** Western blot showing the relative levels of PNKP and PFKFB3 in the nuclear extract of HD patients vs. age-matched control subjects’ frontal cortex. HDAC2: nuclear loading control. **Lower panel:** Quantitation of the relative PFKFB3 levels after normalization with loading control HDAC2. **C. Upper panel:** Representative gel images (four different age-groups representing both genders) showing the 3’-phosphatase activity of PNKP in the post-mortem brain (frontal cortex) nuclear extract of healthy normal control (lane 2) or supplemented with F2,6BP (lane 3, 50 µM), and age/gender-matched HD patient (lane 4) or supplemented with F2,6BP (lanes 5-7, 10, 25 and 50 µM) or F6P (lane 8, 50 µM) or F1,6BP (lane 9, 50 µM). Lane 1: substrate only. Lane 10: purified PNKP (1-2 ng). S: Substrate and P: Released phosphate. **Lower panel:** Quantitation of the % released phosphate in the indicated lanes. The difference between activity between lane 4 (HD patients without F2,6BP supplementation) and lanes 6,7 (with F2,6BP supplementation) is significant (n=3, ***P<0.005). **D.** Bar diagram showing the relative levels of F2,6BP in the nuclear extract of control vs. SCA3 patients’ cerebellum (n=4). **E. Upper panel:** Western blot showing the relative levels of PFKFB3 in the nuclear extract of SCA3 patients vs. age-matched control subjects’ cerebellum (n=3). HDAC2: nuclear loading control. **Lower panel:** Quantitation of the relative PFKFB3 levels after normalization with loading control HDAC2. **F. Upper panel:** Representative gel image (different age-groups, n=4) showing the 3’-phosphatase activity of PNKP in the post-mortem brain (cerebellum) nuclear extract of healthy normal control (lane 2) or supplemented with F2,6BP (lanes 3-4, 25 and 50 µM), and age-matched SCA3 patients (lane 5) or supplemented with F2,6BP (lanes 6-7, 25 and 50 µM) or F6P (lane 8, 50 µM) or F1,6BP (lane 9, 50 µM). Lane 1: substrate only. Lane 10: purified PNKP (2 ng). Lower panel: Quantitation of the % released phosphate in the indicated lanes. The difference between activity between lane 5 (SCA3 patients without F2,6BP supplementation) and lane 7 (with F2,6BP supplementation) is significant (n=3, ***P<0.005). In all the above cases, error bars show ±SD of the mean; n=3 or 4, **P<0.01; ***P<0.005, as applicable.

Next, we tested the effect of F2,6BP supplementation in the restoration of PNKP activity in the nuclear extracts of the patient samples exhibiting decreased F2,6BP levels. Exogenous F2,6BP indeed restored the 3’-phosphatase activity of PNKP in a dose-dependent manner in the nuclear extracts of representative post-mortem HD brains (frontal cortex) across different age and gender groups (**Fig. 3C, lanes 5-7 vs. lane 4**). As controls, we utilized the other two related natural metabolites, F6P and F1,6BP; however, they failed to restore PNKP activity (**Fig. 3C, lanes 8 and 9**). This observation indicates that the specific reduction in F2,6BP levels due to the depletion of PFKFB3 led to the abrogation of PNKP activity and subsequent accumulation of double-strand breaks (DSBs), as observed in our earlier study^27^.

We extended our study to measure F2,6BP levels in the nuclear extracts of post-mortem SCA3 patients (cerebellum) vs. age-matched control subjects (same set as we reported earlier^19^). F2,6BP levels were found to be significantly lower (∼2-fold) in patient samples than that of control **(Fig. 3D)**. PFKFB3 levels were also found to be decreased in the SCA3 patient brain samples compared to age-matched controls (**Fig. 3E**). Notably, exogenous F2,6BP, but neither F6P **(Fig. 3F, lane 8)** nor F1,6BP **(Fig. 3F, lane 9)**, restored the 3’-phosphatase activity of PNKP in a dose-dependent manner **(Fig. 3F, lanes 6-7 vs. lane 5)**. These observations suggest a potential mechanism of vulnerability for a subset of neurons in specific brain regions through the reduction of PFKFB3 and F2,6BP levels. We postulate that under normal physiological conditions, PFKFB3 will increase the local concentration of the metabolite in the nucleus, thus enhancing its availability to PNKP. Alternatively, PFKFB3 hands off F2,6BP to PNKP via their proximity to potentiate DNA repair. Because PFKFB3 is degraded under pathogenic condition and is thus no longer available to hand off or enhance the availability of F2,6BP to PNKP, it may explain a significant decrease in PNKP activity in HD and SCA3.

### Intracellular delivery of F2,6BP can restore PNKP activity and DNA repair deficiency in HD mouse-striatum derived neuronal cells

We next investigated the potential of exogenous F2,6BP in restoring DNA repair deficiency in living cells deficient in PFKFB3. To perform the *in-cell* rescue experiment, we first used HEK293 cells as a model system where PFKFB3 was depleted by siRNA **(Supplementary Fig. S5A)**. We observed impaired DSB repair, evident from widespread accumulation of γH2AX and p53BP1 (markers of DSB accumulation, with p53BP1, specifically indicating impaired C-NHEJ^19,20^) **(Supplementary Fig. S5A, lane 7 vs. lane 1)**. We induced DSBs by bleomycin and allowed cells to repair for 6 h (a time point when efficient repair takes place following DSB induction as tested earlier^18^). Twenty-four hours prior to DSB induction, cells were incubated with F2,6BP in the presence of a cell permeable carrier peptide (K16ApoE; 25 µM)^31^ to facilitate its entry into the cells. Notably, the activation of 3’-phosphatase activity of PNKP was observed exclusively in the nuclear extracts of F2,6BP transfected cells **(Supplementary Fig. S5B, lane 6 vs. lane 2)**, but not in cells transfected with the carrier peptide alone or with F1,6BP **(Supplementary Fig. S5B, lanes 5 and 7 vs. lane 2)**. We also observed a significant reduction in γH2AX and p53BP1 levels only in F2,6BP transfected cells **(Supplementary Fig. S5A, lane 11 vs. lane 9)** whereas the elevated levels persisted in F1,6BP or carrier peptide-treated PFKFB3-depleted cells **(Supplementary Fig. S5A, lane 12 and 10 vs. lane 9)**. These findings strongly indicate that exogenous F2,6BP activates PNKP to potentiate DSB repair in PFKFB3-deficient cells.

Following initial optimization in HEK293 cells, we assessed *in-cell* rescue in striatal neuronal cells derived from Huntington’s disease (HD) mice with expanded polyQ repeats (Q-111) in comparison to wild-type (WT) cells (Q-7)^38^. Biochemical analyses revealed a substantial reduction in the 3’-phosphatase activity of PNKP, but not its protein level^39^, in Q-111 cells (**Fig. 4A, lane 4 vs. lane 2**) correlating with a reduction in PFKFB3 level **(Fig. 4B)** and the subsequent accumulation of DNA strand breaks specifically in the transcribed genome (**Fig. 4C and D, lane 3 vs. lane 1**) compared to Q-7 cells, typical characteristics of the disease pathology. As described above, F2,6BP was delivered with the non-covalent carrier peptide and allowed to alleviate cellular toxicity for 48 and 72 hours. Control experiments involved transfecting Q-111 cells with the carrier peptide alone or with F1,6BP. Importantly, transfection of the peptide alone had no significant toxic effect on PNKP activity in either Q-7 or Q-111 cells (**Fig. 4A, lane 3 vs. lane 2 and lane 5 vs. lane 4**). The restoration of 3’-phosphatase activity of PNKP was observed in a time-dependent manner exclusively in the nuclear extracts of F2,6BP-transfected cells (**Fig. 4A, lanes 6-7 vs. lane 4**), but not in cells transfected with either F1,6BP or the carrier peptide (**Fig. 4A, lanes 8-9 or lane 5 vs. lane 4**). Given that intracellular delivery of F2,6BP restored the 3’-phosphatase activity of PNKP in Q-111 cells, a cell viability assay was performed by crystal-violet staining of live Q-7 and Q-111 cells treated with F2,6BP. Microscopic imaging showed a significant increase in the Q-111 cell count following F2,6BP treatment, comparable to control Q-7 cells, contrary to the decreased count under mock treatment (carrier peptide alone) conditions **(Fig. 4E-G)**. These results collectively suggest that F2,6BP can effectively restore PNKP activity in HD cells, protecting them from neurotoxicity and apoptosis.

**Fig. 4:**
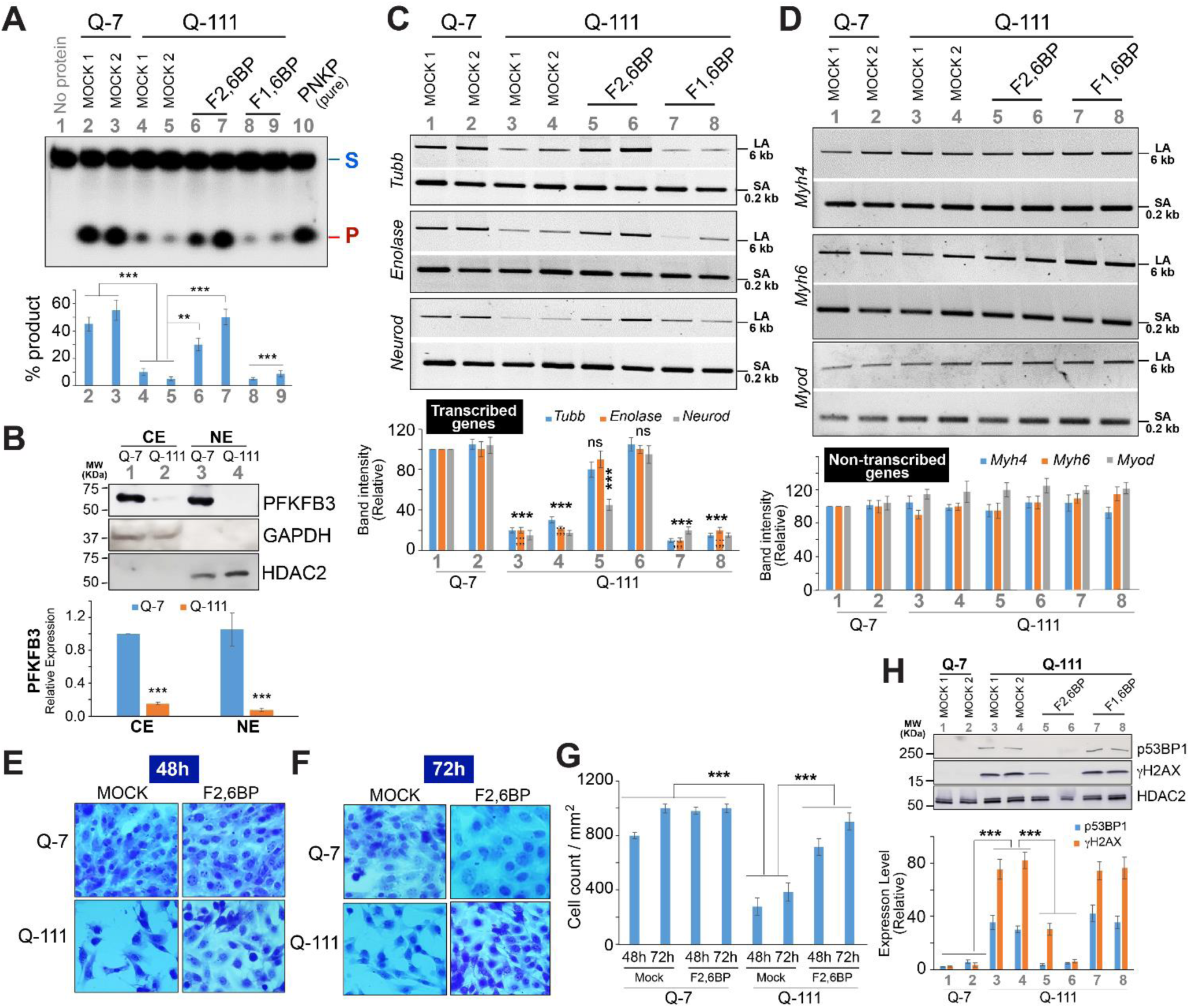
F2,6BP mediated *in cell* rescue of the PNKP activity and TC-NHEJ repair in HD mice striatum-derived neuronal cells (Q-111) **A. Upper panel:** Representative gel image of 3’-phosphatase activity of PNKP in the nuclear extract of Q-7 (lanes 2-3) and Q-111 (lanes 4-9) mice striatum-derived neuronal cells with mock treatment (Mock 1; lanes 2 and 4), treatment with K16ApoE carrier peptide alone (25 µM; Mock 2; lanes 3 and 5) or supplemented with F2,6BP (200 µM; 48 and 72 h) (lanes 6-7), or F1,6BP (lanes 8-9, 200 µM) in presence of carrier peptide. Lane 1: substrate only. Lane 10: purified PNKP (2 ng). **Lower panel:** Quantitation of the % released phosphate in the indicated lanes. Error bars show ±SD of the mean; n=3, **P<0.01; ***P<0.005 between groups as indicated or compared to lanes 2, 3. **B. Upper panel:** The Western blot shows the levels of PFKFB3 in the cytosolic (CE) and nuclear extracts (NE) of Q-7 and Q-111 cells. GAPDH: cytosolic loading control; HDAC2: used as nuclear loading control. **Lower panel:** Quantitation of the relative PFKFB3 levels after normalization with respective loading controls; n=3, ***P<0.005. **C. Upper panel:** Amplification of each long amplicon (6-8 kb) and a small amplicon (∼200-300 bp) of the transcribed (*Tubb*, *Enolase*, *Neurod*) genes to assess DNA strand break accumulation. **Lower panel:** The bar diagram represents the normalized (with short amplicon) relative band intensity with the mock-treated Q-7 sample arbitrarily set as 100. **D. Upper panel:** Amplification of each long amplicon (6-8 kb) and a small amplicon (∼200-300 bp) of the non-transcribed (*Myh4*, *Myh6* and *Myod*) genes to assess DNA strand break accumulation. **Lower panel:** The bar diagram represents the normalized (with short amplicon) relative band intensity with the mock-treated Q-7 sample arbitrarily set as 100. For both C and D, Error bars show ±SD of the mean; n=3, ***P<0.005; ns P>0.05 compared to lanes 1, 2. **E.** Crystal violet (CV)-stained Q-7 **(Upper panel)** and Q-111 **(Lower panel)** cells following mock treatment with carrier peptide **(Left panels)** and supplemented with F2,6BP (200 µM) in the presence of carrier peptide **(Right panels)** at 48 h post F2,6BP delivery. **F.** Similar Crystal violet staining of Q-7 and Q-111 cells at 72 h post F2,6BP delivery. **G.** The bar diagram shows the quantification of cell viability represented as CV-stained cell number/sq. mm area selected from five representative microscopic images of different fields captured independently. Error bars show ±SD of the mean; n=3, ***P<0.005 between indicated groups. **H. Left panel:** The Western blots show the levels of various proteins (indicated on the right) in the nuclear extracts of Q-7 and Q-111 cells under indicated treatment conditions. HDAC2: used as nuclear loading control. **Right panel:** Quantitation of the relative p53BP1 and γH2AX levels after normalization with nuclear loading control HDAC2; n=3, ***P<0.005 between indicated groups.

We then assessed the level of DNA strand breaks in control and F2,6BP-transfected Q-111 or Q-7 cells by LA-qPCR targeting neuron-specific transcribed genes (*Tubb*, *Enolase* and *Neurod*) vs. non-transcribed genes (muscle-specific *Myh4*, *Myh6* and *Myod*). As expected, significant repair of DNA strand breaks was observed only in F2,6BP transfected Q-111 cells **(Fig. 4C, lanes 5-6 vs. lane 3)**. Conversely, F2,6BP showed no notable impact on the repair of non-transcribing genes, as they did not accumulate DNA strand-breaks in Q-111 cells **(Fig. 4D, lanes 5-6 vs. lane 3)**. These results underscore the ability of exogenous F2,6BP to effectively restore PNKP-mediated TC-NHEJ repair in HD cells. We also assessed the repair of DNA strand breaks in PFKFB3-depleted HEK293 cells described earlier by LA-qPCR. Consistent with the results in the neuronal cells, only in F2,6BP transfected cells, but not in the cells treated with either peptide or F1,6BP, significant LA-PCR product was visible indicating efficient DNA strand break repair **(Supplementary Fig. S5C, lane 11 vs. lanes 10 and 12)**.

To further assess the persistence of DNA DSBs or their repair in Q-111 cells under conditions described above, we examined the levels of p53BP1 and γH2AX proteins. In F2,6BP-transfected Q-111 cells, γH2AX and p53BP1 levels were markedly reduced **(Fig. 4H, lanes 5-6 vs. lanes 3),** whereas the elevated levels of γH2AX/p53BP1 (in comparison to Q-7 cells) persisted in either mock or F1,6BP-treated Q-111 cells **(Fig. 4H, lane 4 or lanes 7-8 vs. lane 3)**. These results were consistent with the observations in PFKFB3-depleted HEK293 cells **(Supplementary Fig. S5A)** and demonstrate that F2,6BP deficiency impaired DNA strand break repair, irrespective of cell type specificity.

### F2,6BP can reverse the impaired motor phenotype reported in *Htt*128Q model of Huntington’s disease in Drosophila

To investigate the effect of F2,6BP *in vivo*, we used a well-studied transgenic HD model of the fruit fly (*Drosophila melanogaster*). In this model, the fruit flies were genetically modified to express human HTT gene with an expanded 128 polyQ repeat (*Htt*128Q) under upstream activating sequence (UAS) (BDSC# 33808). This model exhibited the common characteristics of HD pathogenesis, viz., increase in neurotransmitter release, neurodegeneration, reduced lifespan, and impaired motor function^33^. These flies were crossed with specific driver lines (*repo*-GAL4, BDSC# 7415 and *elav*-GAL4, BDSC# 458) for either pan-glial or pan-neuronal expression of *Htt*128Q, respectively **(Fig. 5A, Upper panel)**. The pan-neuronal expression of *Htt*128Q (33808 x 458) showed significant impairment of motor neuron function compared to its pan-glial expression **(Fig. 5A, Lower panel)**. The flies were then supplemented with 1-2 µl of either 200 µM F2,6BP or mock buffer (as control) for 21 days and the rescue of motor deficiency was tested by climbing assays. While there was no detectable effect of this metabolite supplementation in flies expressing *Htt*128Q in glial cells, the impaired motor phenotype was rescued in neuronal cells and a significant increase in climbing score (in both male and female flies) was observed after supplementation with F2,6BP compared to treatment with mock buffer **(Fig. 5A, Lower panel)**. To further confirm whether F2,6BP supplementation can ameliorate repair deficiency, we examined the DNA strand break accumulation in Drosophila CrebB and Neurexin genes by LA-qPCR^19^ in the affected pan-neuronal model. We indeed observed elevated level of DNA damage following *Htt*128Q expression compared to the flies used to generate the strains (*w*^1118^) **(Fig. 5B, lanes 2 and 4 vs. lane 1).** This further indicates that the mock treatment with control buffer failed to rescue genome integrity. However, significant repair was observed following F2,6BP supplementation **(Fig. 5B, lanes 3 and 5 vs. lanes 2 and 4)** and the genome integrity was comparable to the *W*^1118^ flies **(Fig. 5B, Upper panel)**. These findings confirm the ability of F2,6BP to specifically reverse the HD neurodegenerative phenotype *in vivo* in neuronal cells. Overall, our results suggest that supplementing F2,6BP could be a promising approach to prevent key aspects of HD progression via maintaining genomic integrity.

**Fig. 5:**
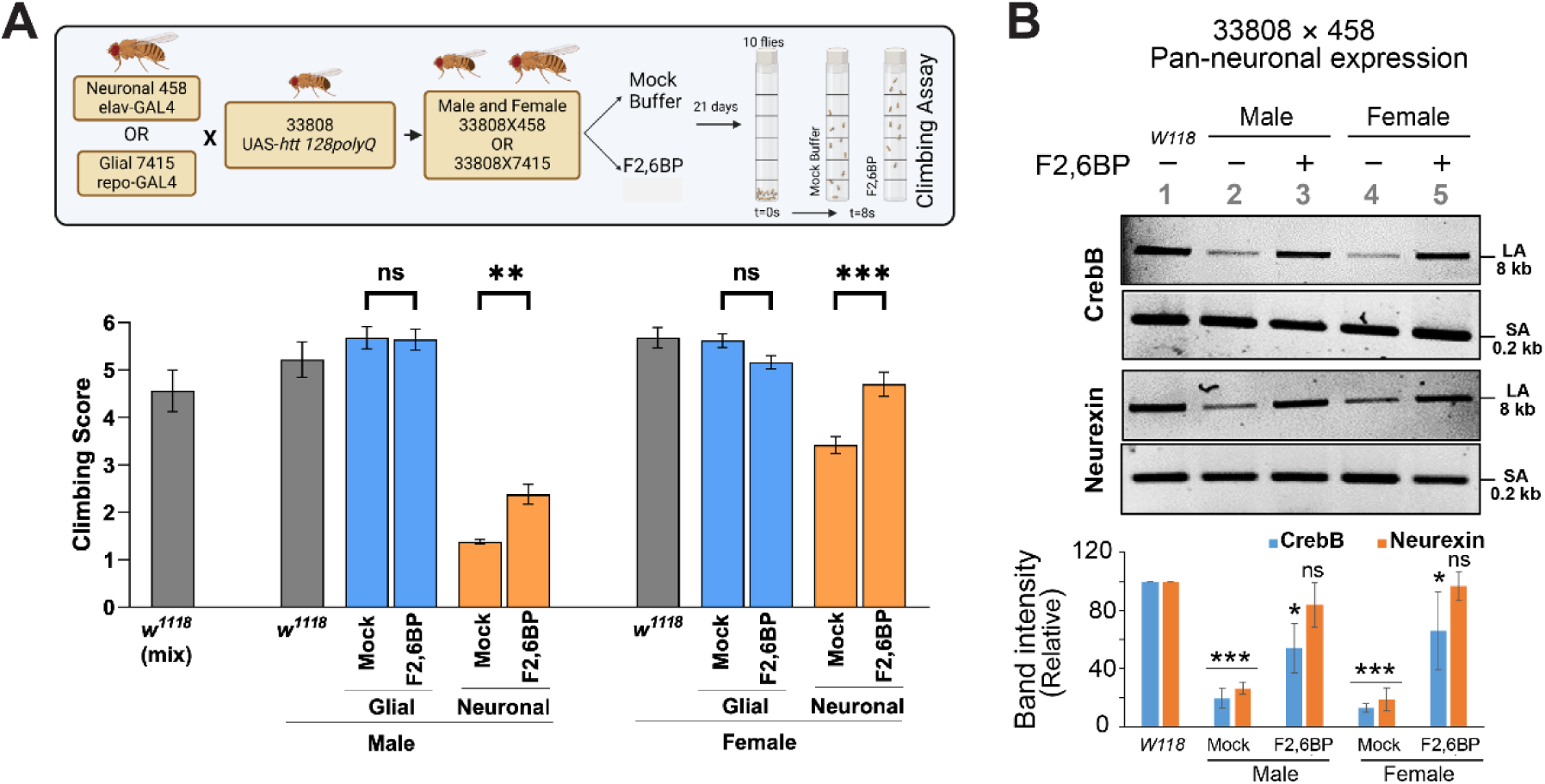
F2,6BP-mediated rescue of neurodegenerative phenotype in a transgenic HD model of Drosophila. **A. Upper panel:** Schematic for climbing score measurement in Drosophila (10 flies) expressing human *HTT* gene with expanded polyQ repeat (*Htt*128Q) in glial and neuronal cells treated with either mock buffer (20 mM Tris-Cl, pH=8.0) or F2,6BP (200 µM) for 21 days and measured at 8 seconds. **Lower panel:** The bar graphs show the effect on climbing score between the mock and F2,6BP treatment or with *w*^1118^ (grey bars) either in male flies or female flies in pan-neuronal (orange bars) or pan-glial (blue bars) expression of *Htt*128Q. Significance (**P<0.01, ***=P<0.005) was assessed using one-way ANOVA uncorrected Fisher’s LSD with a single pooled variance. **B. Upper panel:** Representative agarose gel images of long (∼8 kb; LA) and short (∼200 bp; SA) amplicon of the CrebB and Neurexin genes from genomic DNA of male (lanes 2-3) and female (lanes 4-5) flies with pan-neuronal expression of *Htt*128Q either mock-treated (lanes 2 and 4) or treated with F2,6BP (lanes 3 and 5). Lane 1: *w*^1118^ (males and females). **Lower panel:** The normalized relative band intensities were represented in the bar diagram with *w*^1118^ arbitrarily set as 100 (n=3, error bars represent ±SD of the mean). The damage for each gene in Drosophila with pan-neuronal expression of *Htt*128Q was significant (***=P<0.005) compared to the *w*^1118^ samples. Also, the strand breaks were significantly repaired in F2,6BP-treated samples (*=P<0.05; ns=non-significant P>0.05, compared to *w*^1118^).

The present study uncovers a crucial role of PFKFB3, a nuclear member of the PFKFB family, in transcription coupled DSB repair under normal physiological conditions by maintaining the homeostatic level of F2,6BP, essential for PNKP activity. F2,6BP is traditionally recognized for its role in controlling glycolysis by binding to 6-phosphofructo-1-kinase (PFK1) in the cytoplasm^28^. Our results imply that F2,6BP synthesis by nuclear PFKFB3 is probably a compartmentalized event and locally produced F2,6BP is transferred directly to PNKP within the TC-NHEJ complex. Under the pathogenic HD and SCA3 conditions, PNKP-mediated DNA strand break repair is significantly abrogated due to low levels of nuclear F2,6BP, suggesting a crucial role of F2,6BP *in vivo* for full catalytic activity of PNKP.

The expression of PFKFB3 is stringently controlled by two E3 ligases, anaphase-promoting complex (APC)-Cdh1 (APC^Cdh1^) and beta-transducin repeat-containing protein (β-TRCP). It was reported that under homeostatic conditions, PFKFB3 undergoes constitutive degradation facilitated by APC^Cdh1^ marking a low glycolytic rate and reduced oxidative damage in neurons^40^. However, it has now been shown that PFKFB3 is stabilized during brain excitation to activate glycolysis^41^. This is consistent with our study showing PFKFB3’s level is low only under pathogenic condition but not in post-mitotic healthy neurons. How the expression of polyQ expanded pathogenic proteins, mHTT or mATXN3 reduce PFKFB3 protein levels requires further investigation.

Currently, there is no effective medicine capable of halting or reversing the progression of polyQ disorders. McMurray’s group has earlier developed a synthetic small molecule, XJB-5-131, a radical scavenger and uncoupler of oxidative phosphorylation, that showed promise as a therapeutic compound by suppressing disease phenotypes in HD mouse models^42,43^. Hence, preventing or reversing accumulation of oxidative stress-induced DNA damage under pathogenic condition would be a potential therapeutic option. Here we have demonstrated that exogenous F2,6BP can rescue TC-NHEJ defects and subsequent cellular toxicity in HD cells. Moreover, F2,6BP supplementation rescued neurodegenerative phenotype in a Drosophila model of HD, emphasizing the potential therapeutic efficacy of the endogenous metabolite by protecting brain cells from progressive accumulation of genome damage. To our knowledge, this is the first demonstration of an endogenous metabolite modulating the activity of an essential DNA repair protein. Collectively, our studies thus support the rationale to develop non-toxic analogs of F2,6BP that could specifically target and activate PNKP, benefiting the patients with low levels of F2,6BP.

## Methods

### Cell culture and various treatments

Human Embryonic Kidney 293 (HEK293; ATCC CRL-1573) cells were grown at 37°C and 5% CO2 in DMEM: F-12 (1:1, Cellgro) medium containing 10% fetal bovine serum (Sigma), 100 units/ml penicillin, and 100 units/ml streptomycin (Cellgro). Mouse striatum derived cell line from a knock in transgenic mouse containing homozygous Huntingtin (HTT) loci with a humanized Exon 1 containing 7 or 111 polyglutamine repeats (Q-7 and Q-111; Corriell Institute; Cat# CH00097 and CH00095, respectively) were cultured and maintained in Dulbecco Modified Eagles Medium (high glucose) with 2 mM L-glutamine containing 10% fetal bovine serum, 100 units/ml penicillin and streptomycin, and 0.4 mg/ml G418. As Q-111 cells lose the ability to proliferate and survive, high-passage cultures were avoided. We routinely tested for mycoplasma contamination in cultured cells using the Mycoalert Mycoplasma Detection Kit (Lonza) according to the manufacturer’s protocol, and the cells were found to be free from mycoplasma contamination.

For DNA repair studies, cells were either mock-treated, or treated with bleomycin (Bleo, 30 μg/ml for 1 h; Cat # NDC 0703-3155-01) in reduced serum media (Gibco) and allowed for recovery for 3, 6, 9 and 12 h.

### Analysis of PNKP-associated proteins by 2D gel and MALDI-TOF-TOF MS analysis

A large-scale immunoprecipitation from AGS (human gastric adenocarcinoma) cell nuclear extracts (100 mg, benzonase treated to remove DNA and RNA to avoid DNA-mediated co-immunoprecipitation) were performed using mouse IgG (control) and anti-PNKP antibody (mouse monoclonal Ab, Cytostore)-conjugated agarose beads as described earlier^21^. The immunoprecipitates (IPs) were washed extensively with cold TBS (50 mM Tris-HCl, pH 7.5; 200 mM NaCl) containing 1 mM EDTA, 1% Triton X-100 and 10% glycerol. The complexes were then eluted from the beads stepwise with 25 mM Tris-HCl, pH 7.5 containing 300, 400 and 500 mM NaCl. The eluates were subjected to 2-dimensional gel electrophoresis separation and the protein spots (Sypro Ruby, Molecular Probes) that were specifically present in the PNKP IP and not in the IgG IP were subjected to mass spectroscopic identification in the University of Texas Medical Branch Biomolecular Resource Facility.

### Gene knock-down by siRNA transfection

PFKFB3 depletion was carried out in HEK293 cells using siRNAs (80 nM; transfected twice on consecutive days) from SIGMA (SASI_Hs01_00065120). The cells were treated with control or specific siRNA and lipofectamine 2000 (Invitrogen) mixture for 6 hrs in reduced serum medium (Gibco; reduced serum media) followed by addition of 10% FBS containing DMEM/F12 media for each round of transfection. Nuclear extracts were prepared from the harvested cells (72-96 h post-transfection) to examine the depletion of individual proteins by immunoblot analysis using specific Abs. HDAC2 (Histone deacetylase 2; GTX109642, GeneTex) was used as nuclear loading control.

### Co-immunoprecipitation (Co-IP)

Approximately 100 mg of brain tissue from freshly sacrificed WT mice (6 months old; tissue from three mice were pooled together for a single experiment; n=3) were sliced into small pieces, collected in a pre-chilled, sterile homogenizer (Thomas, PHILA USA C55506) and hand-homogenized with 4 volumes of ice-cold homogenization buffer [0.25 M sucrose, 15 mM Tris-HCl, pH 7.9, 60 mM KCl, 15 mM NaCl, 5 mM EDTA, 1 mM EGTA, 0.15 mM spermine, 0.5 mM spermidine, 1 mM dithiothreitol (DTT), 0.1 mM phenylmethylsulfonyl fluoride (PMSF), and protease inhibitors (EDTA-free; Roche)] with 20 strokes to disrupt tissues^19^. Homogenization was continued until a single-cell slurry was obtained, incubated on ice for 15 min, and centrifuged at 1,000×g to obtain the cell pellet. Nuclear extracts (NEs) were prepared as described^18,19,20^. Briefly, cells were lysed in Buffer A [10 mM Tris-HCl (pH 7.9), 0.34 M sucrose, 3 mM CaCl2, 2 mM magnesium acetate, 0.1 mM EDTA, 1 mM DTT, 0.5% Nonidet P-40 (NP-40) and 1X protease inhibitor cocktail (Roche)] and centrifuged at 3,500×g for 15 min. Nuclear pellets were washed with Buffer A without NP-40 and then lysed in Buffer B [20 mM HEPES (pH 7.9), 3 mM EDTA, 10% glycerol, 150 mM potassium acetate, 1.5 mM MgCl2, 1 mM DTT, 0.1% NP-40, 1 mM sodium orthovanadate (vanadate) and 1X protease inhibitors] by homogenization. Supernatants were collected after centrifugation at 15,000×g for 30 min and DNA/RNA in the suspension was digested with 0.15 U/μl benzonase (Novagen) at 37°C for 1 h. The samples were centrifuged at 20,000×g for 30 min, and the supernatants collected as NEs. Co-IPs were performed using anti-PFKFB3 (GTX108335, GeneTex), Lig IV (Sc-271299, Santa Cruz Biotechnology), and HTT (Sc-477570, Santa Cruz Biotechnology) Abs with Protein A/G PLUS agarose beads (Sc 2003, Santa Cruz Biotechnology) overnight, followed by four washes with Wash buffer [20 mM HEPES (pH 7.9), 150 mM KCl, 0.5 mM EDTA, 10% glycerol, 0.25% Triton-X-100 and 1X protease inhibitors] and eluted with Laemmli Sample Buffer (Bio Rad; final concentration 1X). The immunoprecipitates were tested for the interacting proteins using appropriate Abs [PFKFB3, HTT, ATXN3 (Proteintech 13505-1-AP), PNKP (BB-AB0105, BioBharati Life Science), 53BP1 (Sc-22760, Santa Cruz Biotechnology), DNA-PKcs (GTX6D1 C11 F10, GeneTex), Ku70 (GTX101820, GeneTex), Lig IV (GTX108820, GeneTex), XRCC4 (GTX109632, GeneTex), Polymerase Mu (GTX116332, GeneTex), RNAP II (920202, pSer2, H5 Ab, Biolegend), APE1 (in-house Ab)^19^ and RAD51 (GTX100469, GeneTex)].

### Immunoblotting

The proteins in the nuclear extracts were separated onto a Bio-Rad 4-20% gradient Bis-Tris Gel, then electro-transferred on a nitrocellulose (0.45 μm pore size; GE Healthcare) membrane using 1X Bio-Rad transfer buffer. The membranes were blocked with 5% w/v skimmed milk in TBST buffer (1X Tris-Buffered Saline, 0.1% Tween 20), then immunoblotted with appropriate antibodies [PNKP, PFKFB3, γ-H2AX (S139 residue; #9718S, Cell Signaling Technology), p53BP1 (S1778, #2675S Cell Signaling Technology), 53BP1, HDAC2, H2AX (total) (#2595S; Cell Signaling technology]. The membranes were extensively washed with 1% TBST followed by incubation with anti-isotype secondary antibody (GE Healthcare) conjugated with horseradish peroxidase in 5% skimmed milk at room temperature. Subsequently, the membranes were further washed three times (10 min each) in 1% TBST, developed using ECL^TM^ Western Blotting Detection Reagents (RPN2209, GE Healthcare) and imaged using Kwikquant image analyzer and image analysis software (ver 5.2) (Kindle Biosciences).

### Human tissue samples

Deidentified human post-mortem frontal cortex tissue of HD patients **(Supplementary Table S2)** and cerebellum tissue from SCA3 patients^19^ and age-matched controls (IRB exempt) were obtained from the biorepository of the Michigan Brain Bank, USA through Materials Transfer Agreement (UTMB 22-UFA00474).

### Assay of 3’-phosphatase and 5’-kinase activities of PNKP

The 3’-phosphatase activity of PNKP in the nuclear extract of post-mortem patients’ frontal cortex/cerebellum and age-matched control subjects (2.5 µg) or with purified recombinant PNKP (2 ng) was conducted as we described previously^19,44^. Five pmol of the radiolabeled substrate was incubated at 37°C for 15 min in buffer A (25 mM Tris-HCl, pH 8.0, 100 mM NaCl, 5 mM MgCl2, 1 mM DTT, 10% glycerol and 0.1 μg/μl acetylated BSA). 5 pmol of non-radiolabeled substrate was used as cold substrate. Nuclear extracts were prepared following the protocol used for Co-IP studies. For kinase activity assay, γP^32^ labeled ATP was incubated in kinase assay buffer (80 mM succinic acid pH 5.5, 10 mM MgCl2, 1 mM DTT, 2.5% glycerol) along with 1.0 μg/μl acetylated BSA, and 0.6 pmol labeled substrate for 30 min at 30°C^35^. 100 fmol of PNKP and 2.5 pmole of cold substrate were used in this assay. For *in vitro* PNKP restoration/abrogation, similar assays were done after incubation of F2,6BP/F6P/F1,6BP (in amounts as indicated in the figure legends) with the nuclear extracts for 15 min. The radioactive bands were visualized in PhosphorImager (GE Healthcare) and quantitated using ImageQuant software. The data were represented as % product (released phosphate or kinase product) released from the radiolabeled substrate with a value arbitrarily set at 100%.

### Enzymatic preparation of F2,6BP

Enzymatic preparations of F2,6BP were conducted following the published protocol^45^. Briefly, a reaction cocktail was prepared consisting of 60 mM Tris-HCl (pH 7.5), 1.5 mM DTT, 5 mM Potassium Phosphate (pH 7.5), 20 mM KCl, 40 μM EDTA, 6 mM MgCl2, 5 mM ATP, 1 mM F6P, 10% Glycerol and 1 mg/ml BSA in 200 μL. The reaction was initiated by adding 100 μg PFKFB3 and incubated at 37°C for 90 mins followed by quenching the reaction with 50 μL 1M NaOH and heating at 80°C for 5 mins. The mixture was centrifuged to remove any precipitate followed by dilution to 2 mL with 10 mM Triethylammonium bicarbonate (TEABC) buffer (pH 8.5). The diluted reaction mixture was applied to MonoQ column pre-equilibrated with 10 mM TEABC (pH 8.5). F2,6BP was eluted using 20 to 35% gradient with 800 mM TEABC as buffer B. Peak fractions were pooled based on the phosphate release assay by PNKP, dried and dissolved in 20 mM Tris-Cl (pH=8.0). The presence of F2,6BP was verified by ESI-MS.

### Estimation of F2,6BP in patients’ extracts

To quantify F2,6BP, the F6P assay kit (Sigma-Aldrich) was used to measure the amount of F6P before and after treating the crude cell extracts with 0.1M HCl and incubated at 37°C for 30 mins. Acid treatment converts F2,6BP into F6P. The assay was performed using manufacturer’s protocol which uses NADH-linked luciferase bioluminescence^46^.

### Long amplicon quantitative PCR (LA-qPCR)

The cells were mock- or Bleo-treated 72 h post PFKFB3 depletion and either harvested immediately after Bleo treatment or kept for recovery (3-12 h) after the Bleo treatment and then harvested. Genomic DNA was extracted using the Genomic tip 20/G kit (Qiagen) per the manufacturer’s protocol, to ensure minimal DNA oxidation during the isolation steps. The DNA was quantitated by Pico Green (Molecular Probes) in a black-bottomed 96-well plate and gene-specific LA qPCR assays were performed as described earlier^18,19,20,27,47,48^ using Long Amp Taq DNA Polymerase (New England BioLabs). Three transcribed (HPRT, POLB, and RNAPII, 10.9, 12.1 and 11.3 kb, respectively) and three non-transcribed (NANOG, 8.6 kb, OCT3/4, 10.1 kb, MyH2, 6.0 kb) genes were amplified from HEK293 cells using appropriate oligos.

Similar LA-qPCR was performed from genomic DNA isolated from Q-7 and Q-111 cells. Since these are neuronal cells, a different set of transcribed (neuronal differentiation factor 1 (*Neurod*), tubulin β3 class III (*Tubb*) and gamma-enolase (*Enolase*) vs. non-transcribed (myogenic differentiation factor [*Myod*], muscle-specific myosin heavy chain 4 and 6 [*Myh4*, *Myh6*]) genes were used for the LA-qPCR assay.

Finally, genomic DNA isolated from adult Drosophila (10 from each genotype and different treatment groups) were used for DNA damage analysis. Two genes (CrebB and Neurexin, ∼8 kb) were amplified using appropriate oligos.

The LA-qPCR reaction was set for all genes from the same stock of diluted genomic DNA sample, to avoid variations in PCR amplification during sample preparation. Preliminary optimization of the assays was performed to ensure the linearity of PCR amplification with respect to the number of cycles and DNA concentration (10-15 ng). The final PCR reaction conditions were optimized at 94°C for 30 s; (94°C for 30 s, 55-60°C for 30 s depending on the oligo annealing temperature, 65°C for 10 min) for 25 cycles; 65°C for 10 min. Since amplification of a small region is independent of DNA damage, a small DNA fragment (∼200-400 bp) from the corresponding gene(s) was also amplified for normalization of amplification of the large fragment. The amplified products were then visualized on gels and quantitated with ImageJ software (NIH). The extent of damage was calculated in terms of relative band intensity with a control siRNA/mock-treated sample or *w*^1118^ sample (for Drosophila studies) considered as 100. All oligos used in this study are listed in **Supplementary Table S3**.

### *In cell* delivery of the exogenous F2,6BP/F1,6BP in Q-7 and Q-111 cells

For delivery of the glycolytic metabolites in the mouse striatum-derived neuronal cells (Q-7 and Q-111), we followed the protocol optimized in our lab^32^. Briefly, 200 μM of F2,6BP or F1,6BP was mixed with 25 μM cell-permeable carrier peptide K16ApoE (Mayo Proteomic Core Facility) and incubated for 45 min at RT; then metabolite-peptide mix was added to the cells and incubated for 6 h in reduced serum media. After 6 h, FBS containing complete media was added and cells were harvested at 48 and 72 h for DNA repair assays/LA-qPCR. In another case, 15,000 cells were plated in 24 well-flat-bottomed plates in 2 ml medium per well. Following mock or F2,6BP delivery in the dose mentioned above, the cells were allowed to grow for additional 48-72 hours, culture medium was aspirated, and cells were stained with 10 μl crystal violet dye solution (to image the live cells). Microscopic imaging was done under bright field using WHN10×/22 eyepiece and a 20× objective (field of view is 1.1 mm, and camera correction is 1.0) on an Echo Revolution Microscope system. More than 5 randomly selected fields of view per sample were photographed and cells were counted manually/sq.mm of the chosen optical field.

Delivery of the glycolytic metabolites, F2,6BP/F1,6BP were performed using a similar protocol in HEK293 cells 48h following depletion of PFKFB3 and 24h prior to treatment with bleomycin.

### Protein expression and purification

WT recombinant His-tagged PFKFB3 and PNKP were purified from *E. Coli* using protocol as described earlier^21,32^. Briefly, pET28a (Novagen) vector containing N-terminal His-tagged-WT PFKFB3 or PNKP coding DNA sequence was transformed into *E. coli* BL21(DE3) RIPL Codon-plus cells. Log-phase culture (A600 = 0.4–0.6) of *E. coli* was induced with 0.5 mM isopropyl-1-thio-β-D-galactopyranoside at 16 °C for 16 h. After centrifugation, the cell pellets were suspended in a lysis buffer (buffer A) containing 25 mM Tris-HCl, pH 7.5, 500 mM NaCl, 10% glycerol, 1 mM β-mercaptoethanol (β-ME), 0.25% Tween 20, 5 mM imidazole, 2 mM PMSF. After sonication, the lysates were spun down at 13,000 rpm, and the supernatant was loaded onto HisPur Cobalt Superflow Agarose (Thermo Scientific, catalog no. 25228) previously equilibrated with buffer A and incubated for 2 h at 4 °C. After washing with buffer A with gradient increasing concentration of imidazole (10, 20, 30, 40 mM), the His-tagged proteins were eluted with an imidazole gradient (80–500 mM imidazole in buffer containing 25 mM Tris-HCl, pH = 7.5, 300 mM NaCl, 10% glycerol, 1 mM ß-ME, 0.25% Tween 20). After elution, the peak protein fractions were dialyzed against buffer C (1XPBS, pH 7.5, 1 mM DTT, and 25% glycerol).

GST-tagged WT PNKP and its domains were purified as described previously^21^, following protocols described as above. Glutathion-sepharose is used as bead instead of HisPur Cobalt Superflow Agarose.

### GST or His pull-down assays for *in vitro* interaction study

GST pulldown assays were performed as described previously^21^. Briefly, GST-tagged full-length PNKP or its three individual domains (20 pmol) were bound to glutathione-Sepharose beads (20 μL), washed thoroughly with buffer A (25 mM Tris-Cl pH 7.5, 0.1% Triton X-100, 0.1 mM EDTA and 10% glycerol) containing 150 mM NaCl, and then incubated with WT PFKFB3 (20 pmol) with constant rocking for 4 h at 4°C in 0.5 ml of 150 mM salt containing buffer A. After extensive washing with 200 mM NaCl containing buffer A, 20% of the bound proteins were separated by SDS-PAGE for immunoblotting analysis using an anti-PFKFB3 or anti-GST Ab.

For His pulldown assay, His-tagged PFKFB3 was bound to HisPur cobalt agarose beads (20 μL), washed in buffer A with 150 mM NaCl and then incubated with GST-tagged PNKP/domains (20 pmol) as described above.

### Drosophila maintenance and treatment

All Drosophila stocks were maintained at 25°C on standard fly food under a 12:12-h light-dark cycle. Drosophila strains were purchased from Bloomington Drosophila Stock Center (BDSC, Bloomington, Indiana, USA: UAS-*htt* (#33808, RRID: BDSC_33808), expresses human Huntingtin (HTT) with long polyQ (glutamine) repeat of 128 amino acids. The repo-GAL4 (#7415, RRID: BDSC_7415) is a pan-glial promoter and *elav*-GAL4 (#458, RRID: BDSC_458) is a pan-neuronal promoter. BDSC 33808 were crossed to BDSC_7415 or BDSC_458 for pan glial or pan-neuronal expression (shown schematically in **Fig. 5A**). Flies eclosed from the crosses were collected in fresh food vials and kept under standard conditions for 1-2 days for acclimatization. Flies were separated into two vials for Mock buffer and F2,6BP treatment. One BD syringe (1 mL) was filled with Mock buffer, and the other was filled with F2,6BP and drop is administered per day to respective cohorts at the same time of the day for 21 consecutive days.

### Climbing or Negative Geotaxis assay

The climbing assay was performed as described with minor modifications^19^. Briefly, experimental flies were anesthetized on ice. A group of 10 male flies per vial were transferred to a 25 mL sterile glass measuring cylinder. The measuring cylinder was divided into 6 compartments equally, the lowest compartment was labeled with 1 and the highest compartment was labeled with 6. The measuring cylinder with flies were placed against a white background for better video recording. The cylinder was tapped gently 3 times to send the flies to the bottom of the cylinder. The climbing time was recorded for 20 seconds. Five trials were performed for each cohort. The climbing score was calculated at 8 seconds.

## Acknowledgements

This work was supported by National Institute of Health Grants 2R01 NS073976 and R56NS073976 to TH, R01AI163327 and R21 AG078635 to GG, Don and Nancy Mafrige Professorship in Neurodegenerative Diseases (BK), Alzheimer’s Association Research Grant (AARG-17-533363 BK), National Institutes of Aging (R21-AG059223 and R01-AG063945 BK), Ministerio de Ciencia e Investigación, AEI 10.13039/501100011033 (PID2020-113238RB-I00), Comunidad de Madrid and CIBERCV to LB. We thank the Michigan Brain Bank (5P30 AG053760 University of Michigan Alzheimer’s Disease Core Center) for providing us with the tissue of post-mortem HD and SCA3 patients and their age-matched controls. We thank Dr. Sankar Mitra for carefully reading this manuscript.

## Author contributions

TH conceived the research. TH, GG, AC and BK designed the research. AC, SGS, WM, WH, TB, and SMM performed research. LB provided reagents and critical intellectual inputs. AC, SGS, BK, TB, GG and TH wrote the manuscript. All the authors read and approved the final version of the manuscript.

## Conflict of interest

The authors declare that they do not have any conflicts of interest.

## Data availability

The data that support the findings of this study are available from the corresponding author upon request.

---

**Supplementary Fig. S1.**
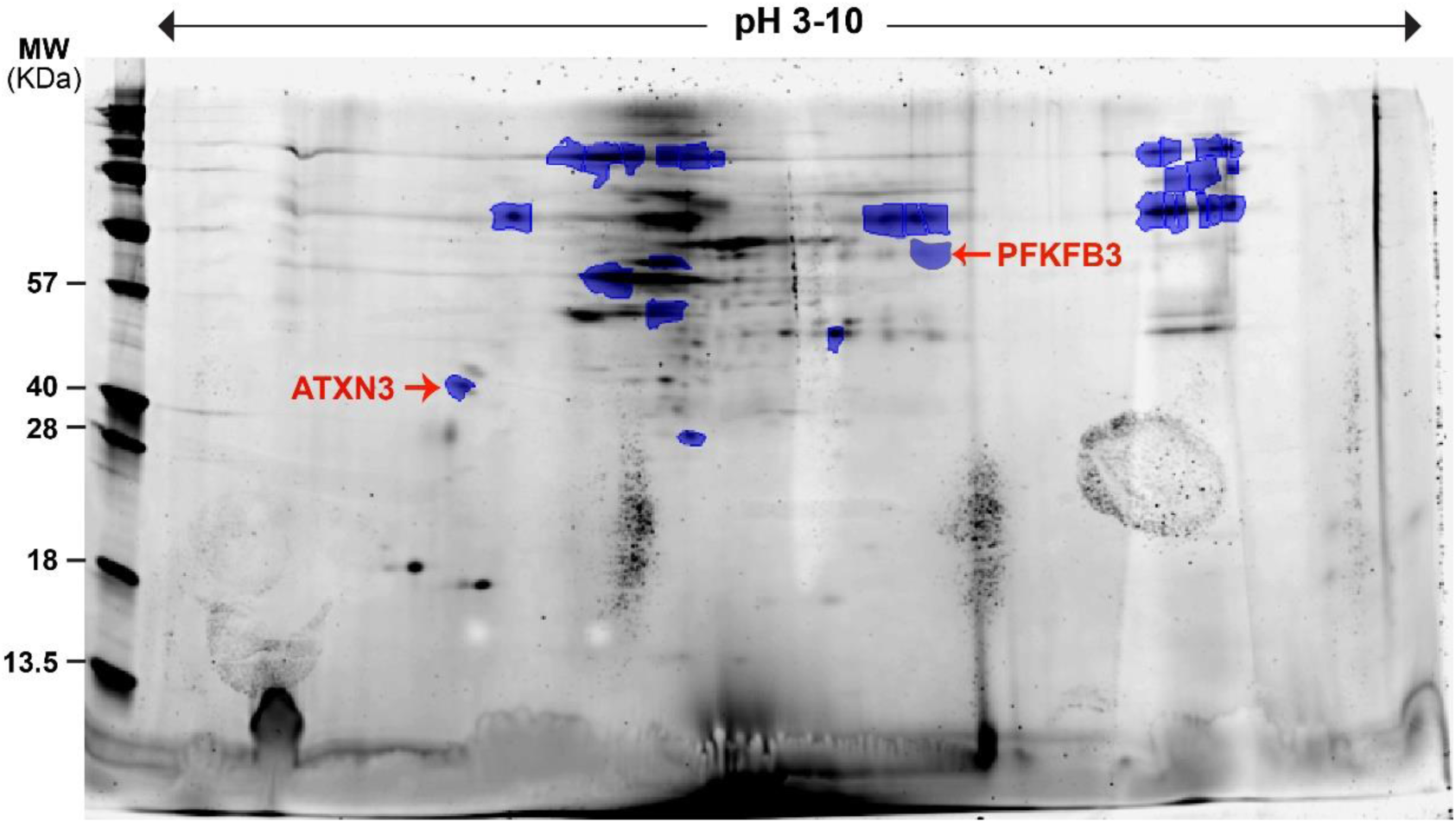
Identification of PFKFB3 in the PNKP IP by 2D gel and MALDI-TOF-TOF MS analysis. The spot annotated was identified as human PFKFB3 by subsequent MS/MS analysis (See Supplementary Table S1). ATXN3 (reported in our previous publication https://doi.org/10.1371/journal.pgen.1004749) is also annotated in the same figure. Image re-used on permission from our earlier published article: Chatterjee A, Saha S, Chakraborty A, Silva-Fernandes A, Mandal SM, Neves-Carvalho A, et al. (2015) The Role of the Mammalian DNA End-processing Enzyme Polynucleotide Kinase 3’-Phosphatase in Spinocerebellar Ataxia Type 3 Pathogenesis. *PLoS Genet* 11(1): e1004749. https://doi.org/10.1371/journal.pgen.1004749.

**Supplementary Fig. S2.**
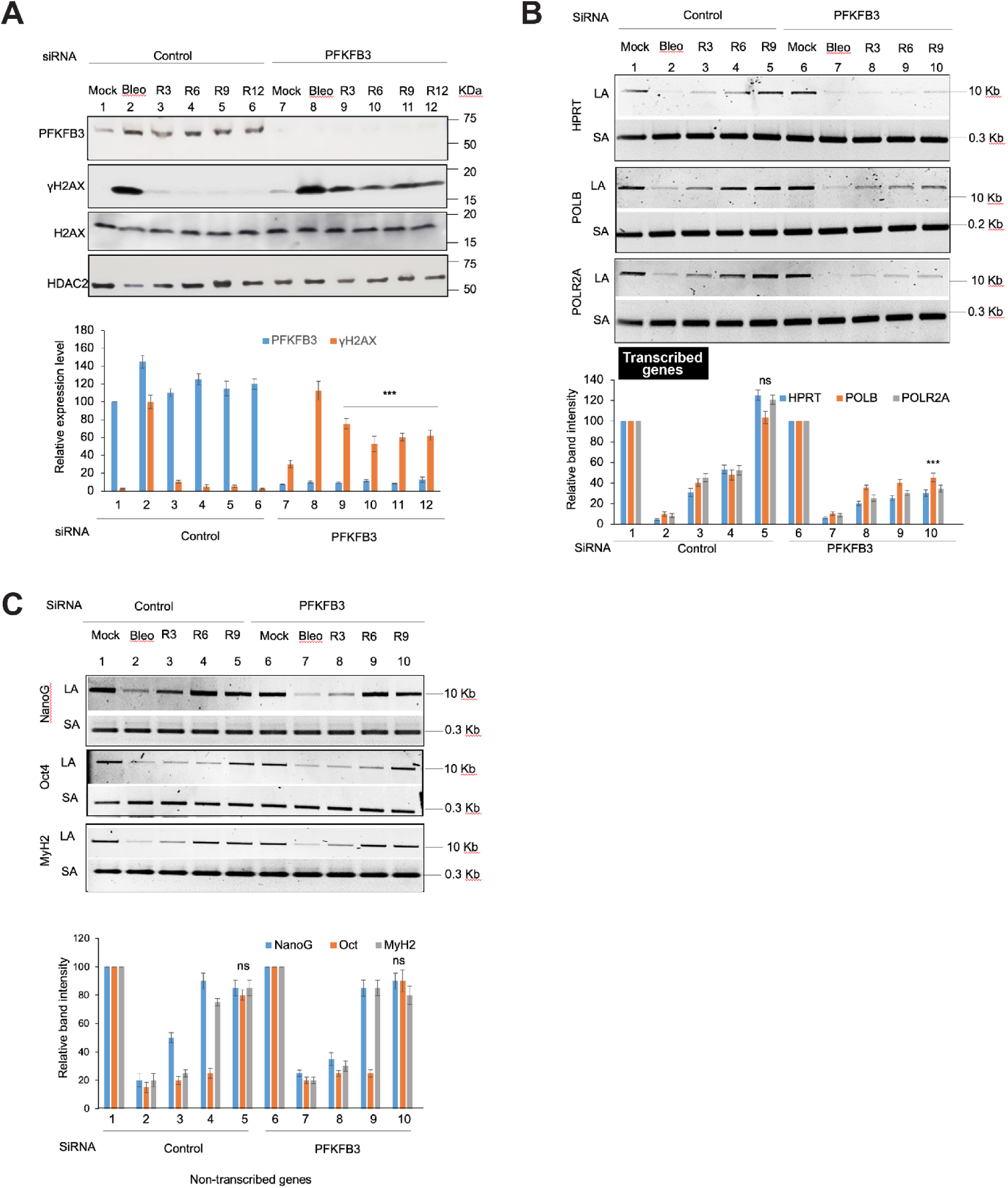
Effect of PFKFB3 depletion on TC-NHEJ repair. **A. Upper panel:** Western blot showing the levels of indicated proteins (on the left) in the nuclear extract of control siRNA (lanes 1-6) and PFKFB3 siRNA (lanes 7-12) transfected HEK293 cells either mock, Bleo-treated or 3 (R3), 6 (R6), 9 (R9) and 12 (R12) h recovery following Bleo treatment. HDAC2: used as nuclear loading control. **Lower panel:** Quantitation of the relative PFKFB3 and γH2AX levels after normalization with nuclear loading control HDAC2; n=3, ***P<0.005 between lanes 3-6 (recovery in control cells) and lanes 9-12 (recovery in PFKFB3 depleted cells). **B.** HEK293 cells were transfected with either control siRNA (lanes 1-5) or PFKFB3 siRNA (lanes 6-10) and further mock- or Bleo-treated or kept for recovery for 3 (R3), 6 (R6) and 9 (R9) h after Bleo treatment. **Upper panel:** Amplification of each long amplicon (LA) (10-12 kb) and a small amplicon (SA) (∼200-300 bp) of the transcribed (HPRT, POLB and POLR2A) genes. **Lower panel:** The bar diagram represents the normalized (with short amplicon) relative band intensity with the mock-treated sample arbitrarily set as 100. **C.** Similar experiment showing amplification of each long amplicon (LA) (10-12 kb) and a small amplicon (SA) (∼200-300 bp) of the non-transcribed (NanoG, Oct4, MyH2) genes **(Upper panel)**. **Lower panel**: The bar diagram represents the normalized relative band intensity with the mock-treated sample arbitrarily set as 100. For both B and C, Error bars show ±SD of the mean; n=3, ***P<0.005 or ns=P>0.05, compared to respective mock-treated sample.

**Supplementary Fig. S3.**
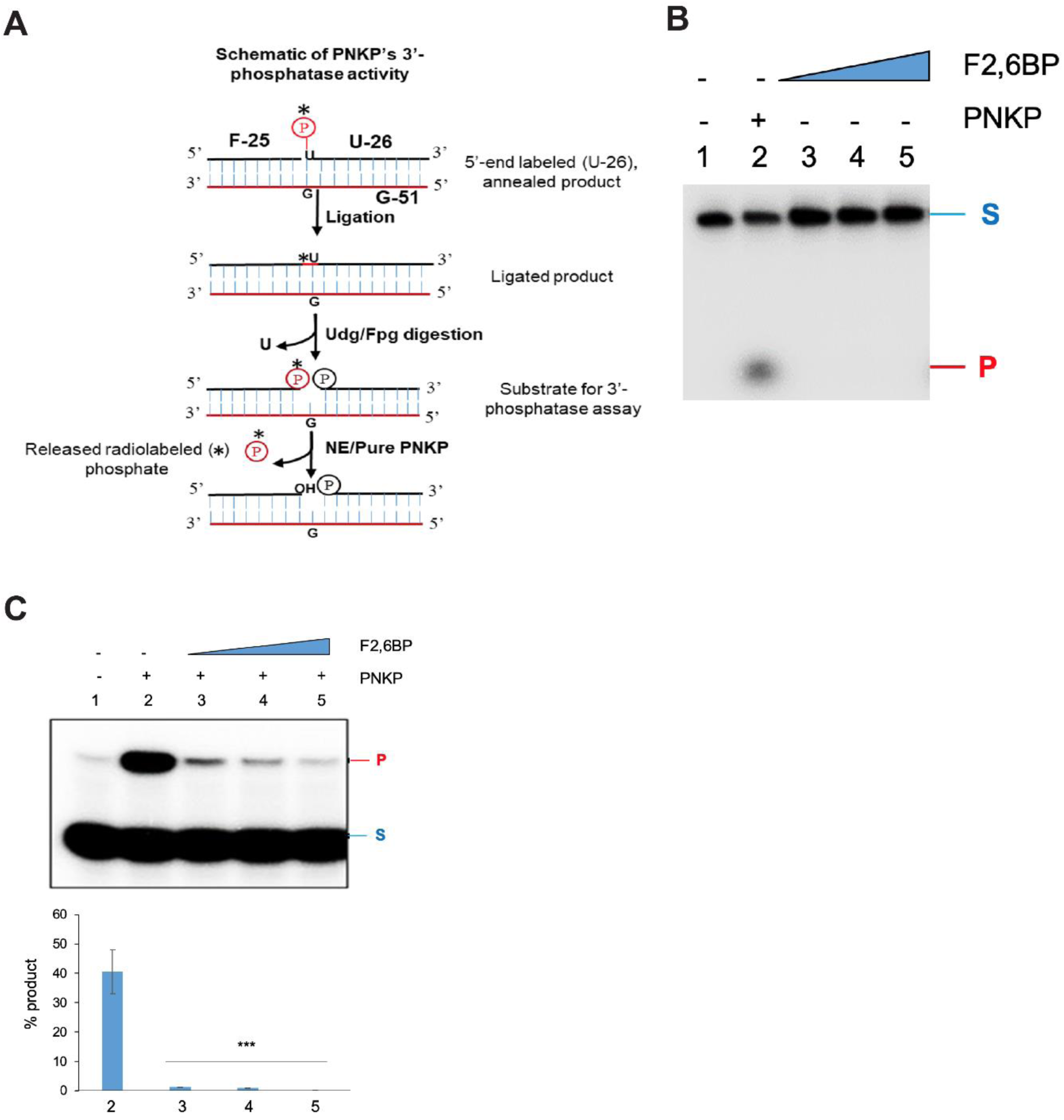
Effect of fructose 2,6-bisphosphate on PNKP activity. **A.** Schematic representation of 3’-phosphatase activity assay of PNKP. **B.** Lane 2: 3’-phosphatase activity of purified PNKP (2 ng). Lane 1: substrate only. Lane 3-5: similar 3’-phosphatase assay with increasing doses of F2,6BP (50 µM-500 µM-5 mM). S: Substrate and P: Released phosphate. **C. Upper panel**: 5’-kinase activity of purified PNKP alone (200 fmole, lane 2) PNKP plus increasing amounts of F2,6BP (10, 15 and 20 µM, lanes 3-5). Lane 1: substrate only. **Lower panel:** Quantitation of the % kinase product. Error bars show ±SD of the mean; n=3, ***P<0.005.

**Supplementary Fig. S4.**
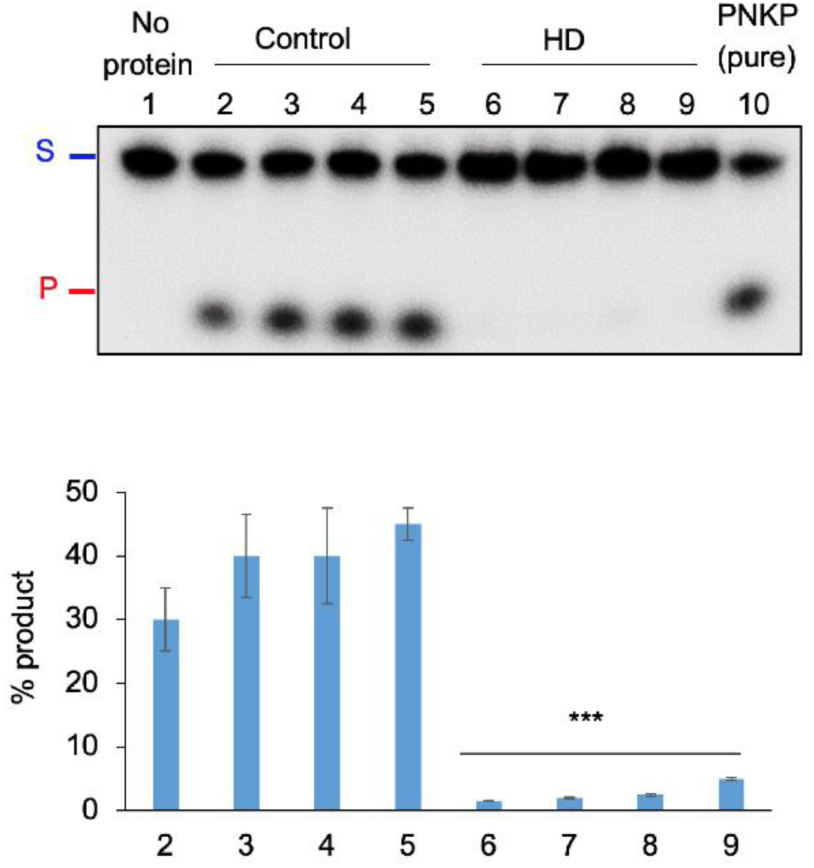
**Upper panel:** 3’-phosphatase activity of PNKP in the nuclear extract (250 ng) of frontal cortex from healthy normal control (male) (lanes 2-5) vs. age-matched HD patients (lanes 6-9). Lane 1: substrate only. Lane 10: Purified PNKP (2 ng). S: Substrate and P: Released phosphate. **Lower panel:** Quantitation of the % released phosphate in the indicated lanes; n=3, ***P<0.005 between Control vs. HD patient.

**Supplementary Fig. S5.**
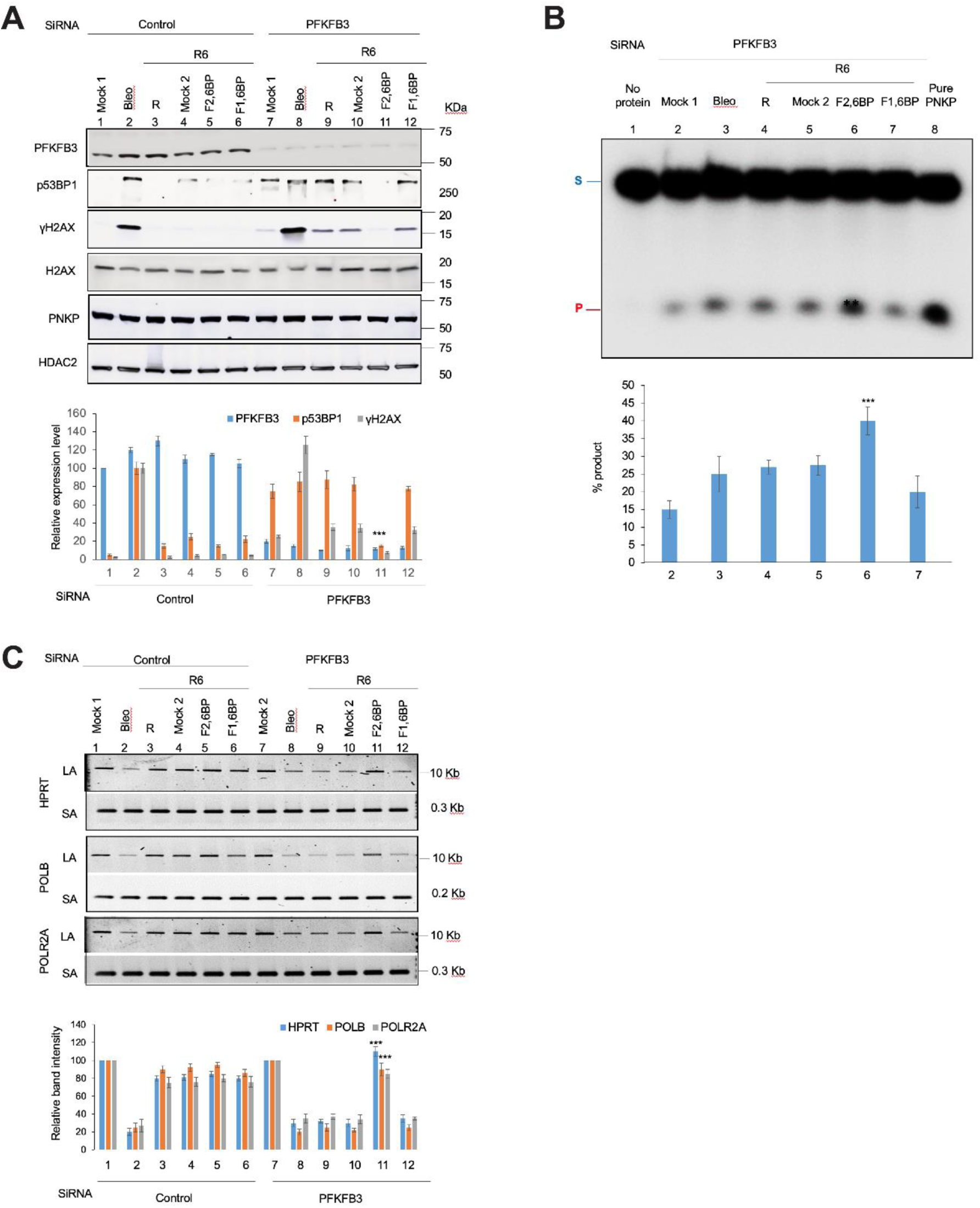
F2,6BP mediated rescue of the PNKP-mediated TC-NHEJ repair in HEK293 cells **A. Upper panel:** The Western blot shows the levels of various proteins (indicated on the left) in the nuclear extracts of control (lanes 1-6) or PFKFB3 siRNA (lanes 7-12) transfected HEK293 cells. Mock 1: Mock treatment; Mock 2: treatment with K16ApoE carrier peptide (25 µM); Bleo: Bleo treatment; R6, R: Recovery for 6 h after Bleo treatment; F2,6BP: Prior treatment with 200 µM F2,6BP + carrier peptide; F1,6BP: Prior treatment with 200 µM F1,6BP + Carrier peptide. HDAC2: used as nuclear loading control. **Lower panel:** Quantitation of the relative PFKFB3, p53BP1 and γH2AX levels after normalization with nuclear loading control HDAC2; n=3, ***P<0.005 for p53BP1 and γH2AX levels between lane 11 and lanes 9, 10, 12. **B. Upper panel**: Representative gel image of 3’-phosphatase activity of PNKP in the nuclear extract of PFKFB3 siRNA transfected cells under similar conditions. Lane 1: substrate only. Lane 8: purified PNKP (2 ng). **Lower panel:** Quantitation of the % released phosphate in the indicated lanes. Error bars show ±SD of the mean; n=3, ***P<0.005 compared to mock treatment (Mock 1). **C. Upper panel:** Amplification of each long amplicon (10-12 kb) and a small amplicon (∼200-300 bp) of the transcribed (HPRT, POLB and POLR2A) genes to assess DNA strand break accumulation. **Lower panel:** The bar diagram represents the normalized relative band intensity with the mock-treated sample arbitrarily set as 100. Error bars show ±SD of the mean; n=3, ***P<0.005 compared to samples in lanes 8, 9, 10 and 12.

**Supplementary Table S1:**
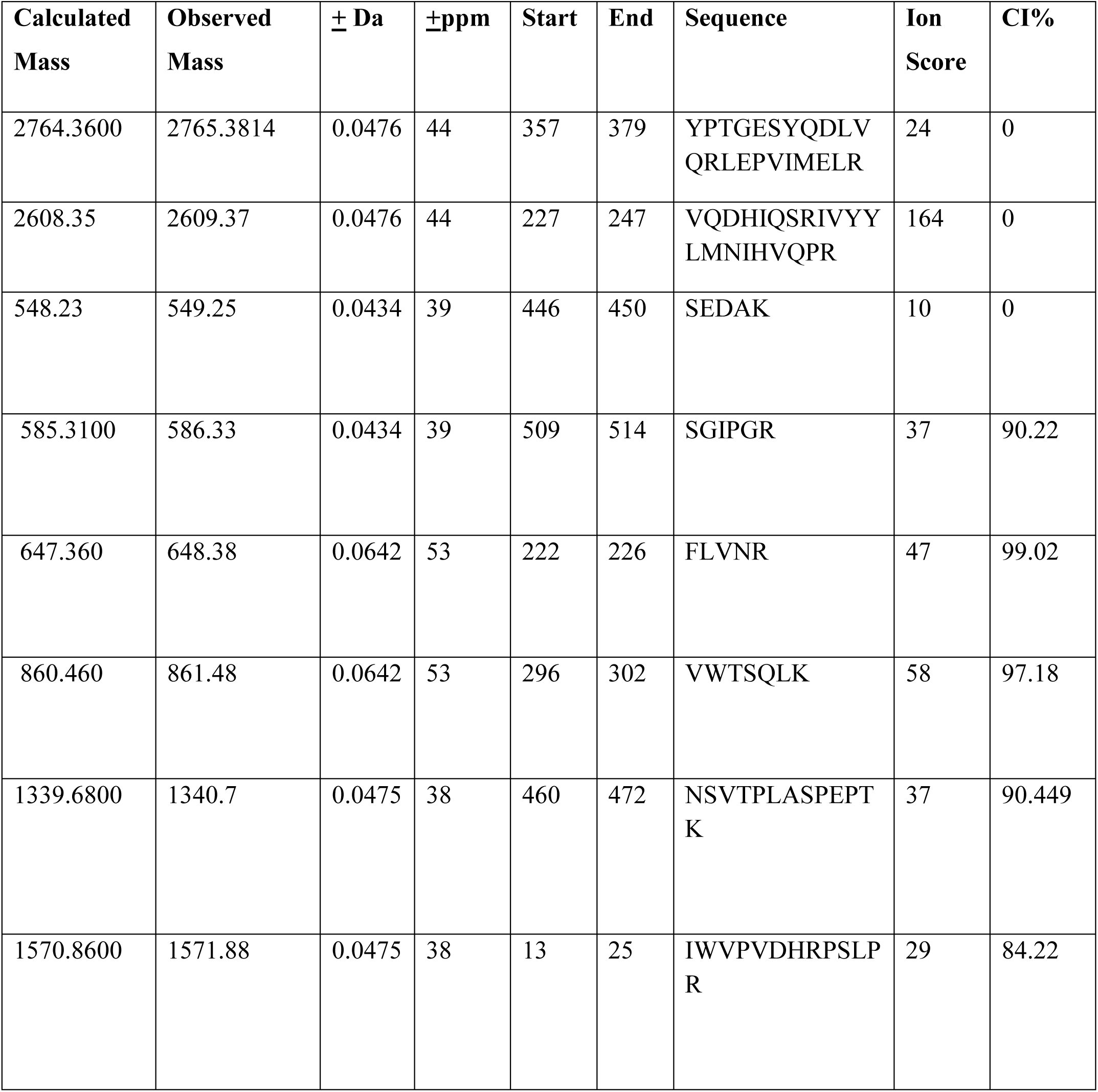
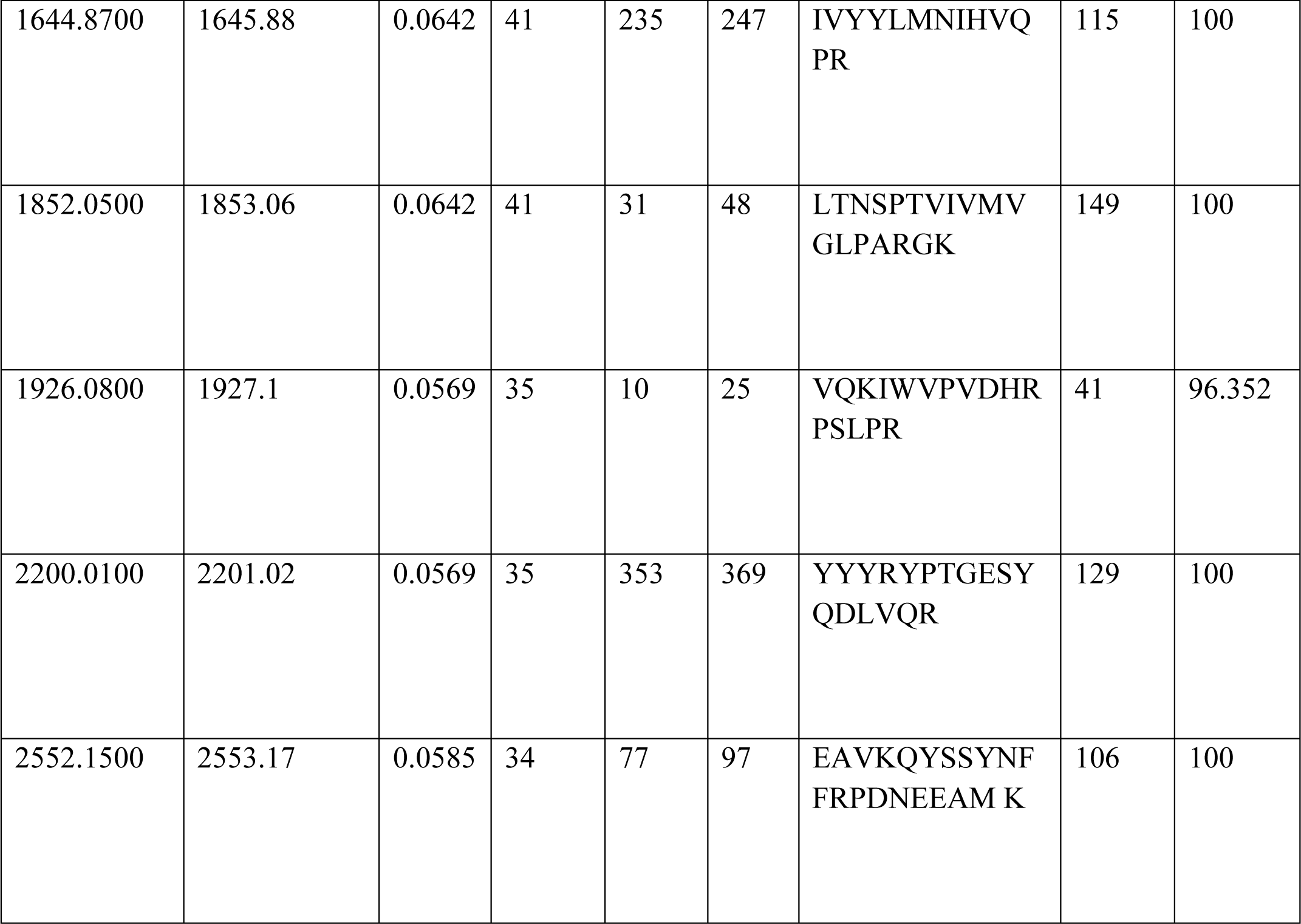
Analysis of mass spectrometry data for PFKFB3-derived peptides.

**Supplementary Table S2:** Details of the human post-mortem brain tissues (frontal cortex)

| Brain Bank ID | Age of Death | Sex | PMI (hours) | Neuropathology |
| --- | --- | --- | --- | --- |
| 115 | 62 | Male | 9 | HD |
| 1333 | 58 | Male | 19 | HD |
| 1174 | 54 | Male | 7 | HD |
| 927 | 53 | Male | 22 | HD |
| 423 | 41 | Male | 7 | HD |
| 761 | 70 | Male | 16 | HD |
| 1122 | 76 | Male | 8 | HD |
| 1024 | 68 | Male | 5 | HD |
| 1017 | 65 | Male | 12 | HD |
| 107 | 76 | Female | 8 | HD |
| 1459 | 72 | Female | 10 | HD |
| 2241 | 68 | Female | 25 | HD |
| 1121 | 67 | Female | 8 | HD |
| 1232 | 61 | Female | 6 | HD |
| 1508 | 56 | Female | Not recorded | HD |
| 1298 | 49 | Female | 13 | HD |
| 1071 | 44 | Female | 7 | HD |
| 14 | 65 | Male | 24 | Control |
| 609 | 61 | Male | 24 | Control |
| 729 | 59 | Male | 12 | Control |
| 260 | 39 | Male | Not recorded | Control |
| 915 | 81 | Male | 14 | Control |
| 283 | 70 | Male | 21 | Control |
| 899 | 76 | Female | 14 | Control |
| 57 | 74 | Female | 6 | Control |
| 1539 | 53 | Female | 11 | Control |

**Supplementary Table S3:**
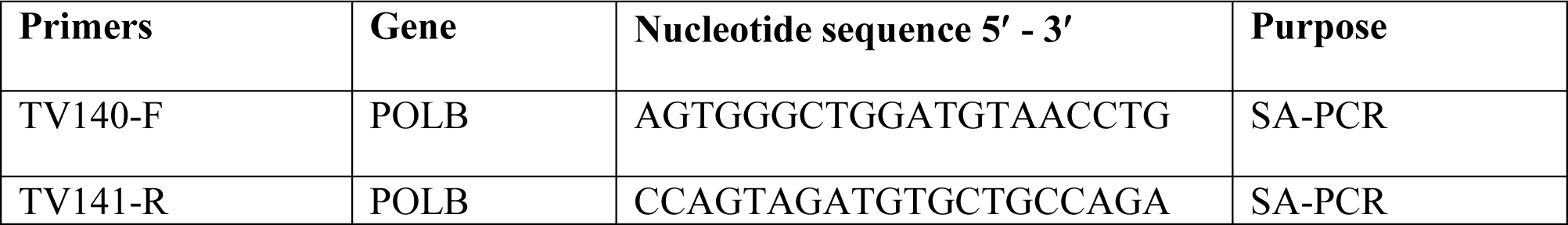
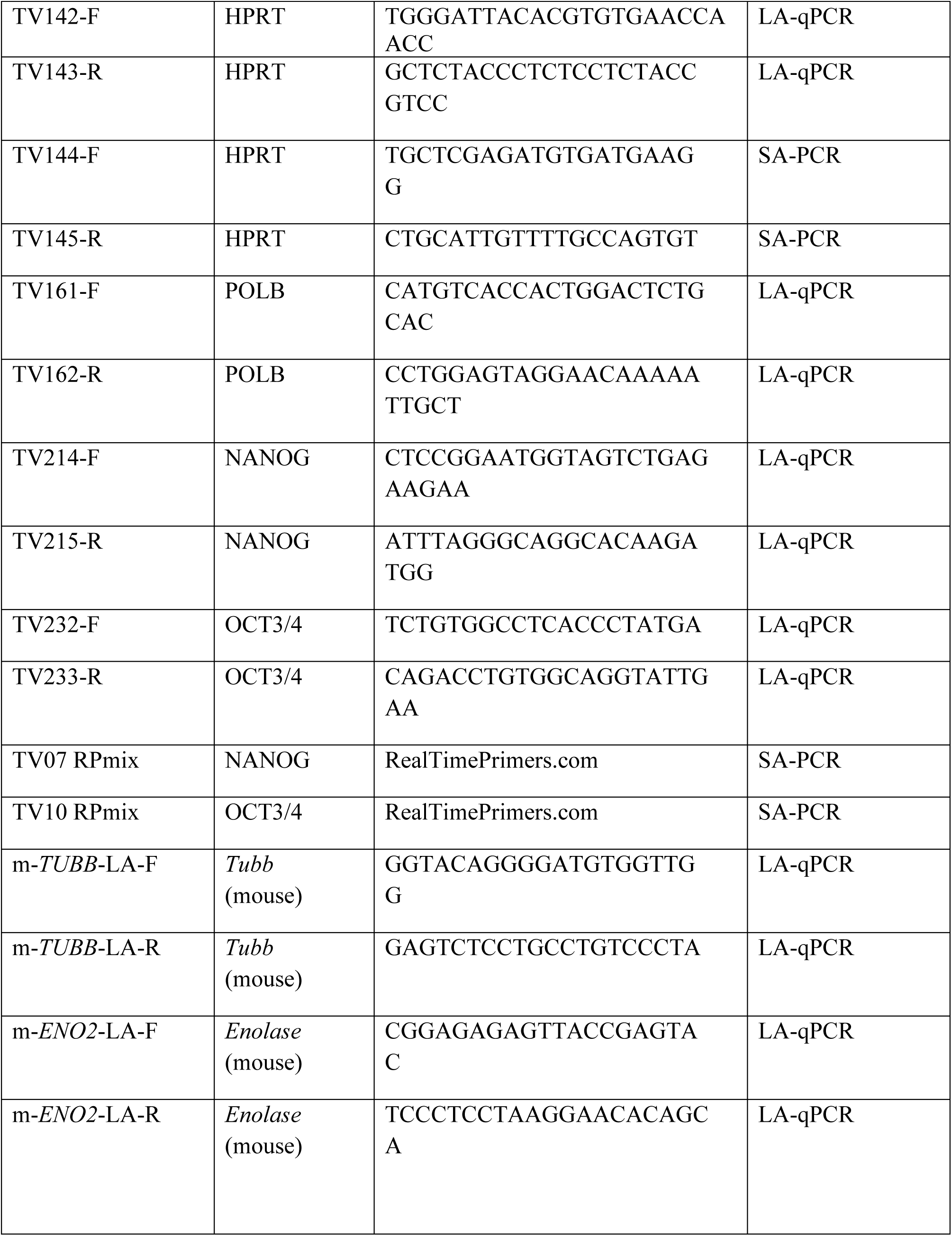
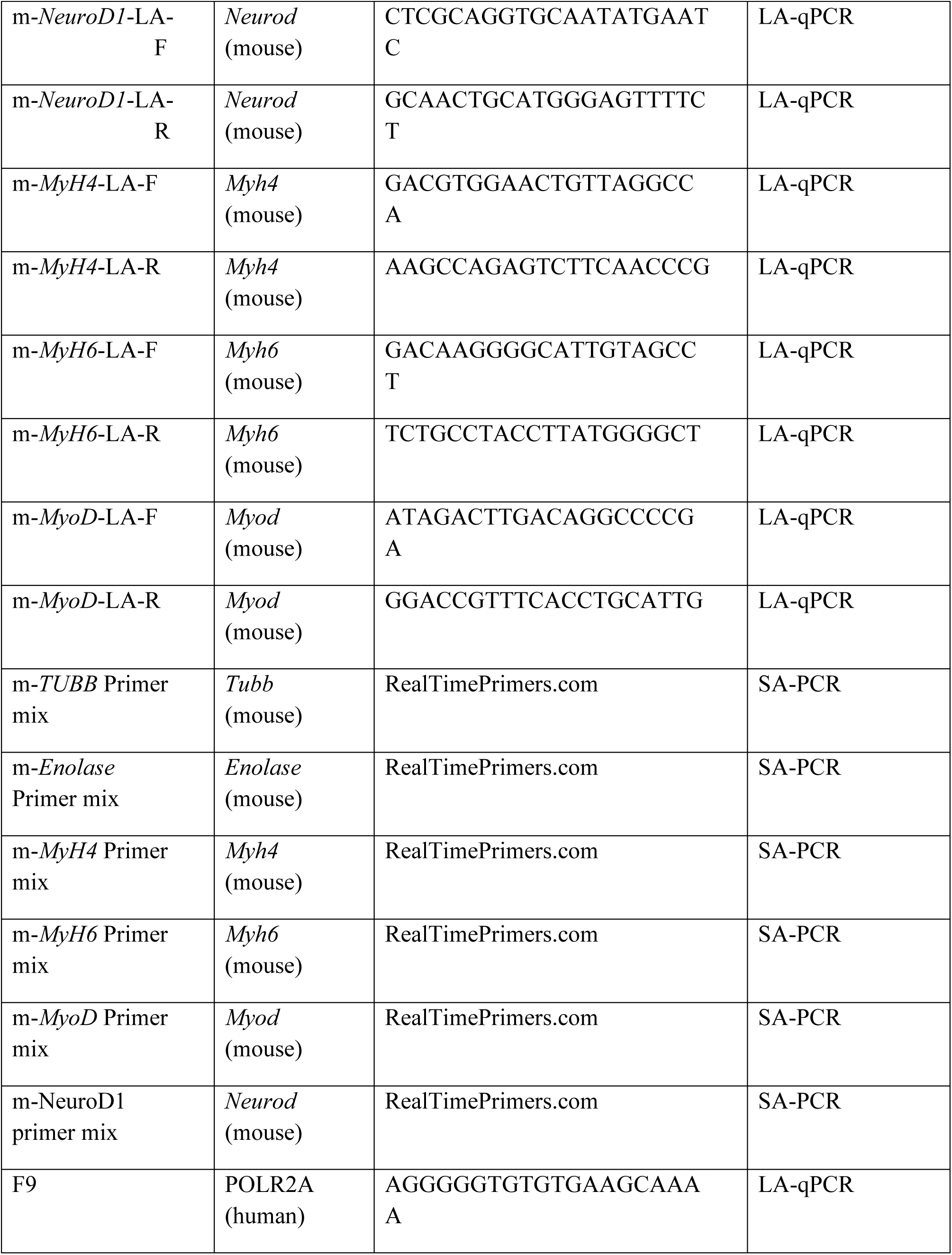
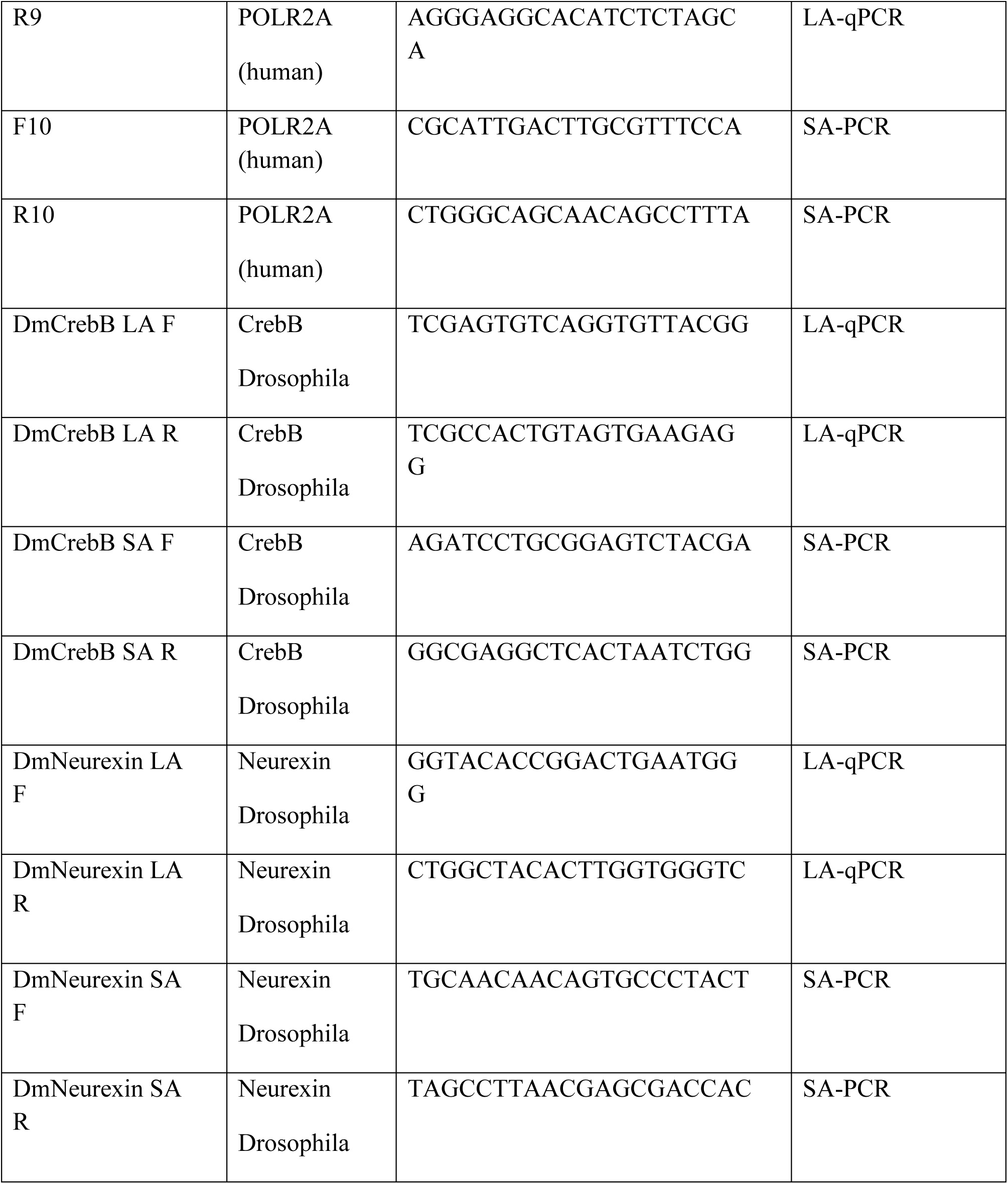
Primers used in the study.

## Notes

### Competing Interest Statement

The authors have declared no competing interest.

### Summary of Updates

We revised the manuscript to incorporate the new results shown in Fig. 1A, 1D, 1E, 4B and Supplementary Fig. S3B and modified the title to Fructose-2,6-bisphosphate restores DNA repair activity of PNKP and ameliorates neurodegenerative symptoms in Huntingtons disease. We believe that the new title adequately described the main message of the article. We also included revised Figures 3C and 3F.

